# A phosphoproteomic approach reveals that PKD3 controls phenylalanine and tyrosine metabolism

**DOI:** 10.1101/2020.07.20.211474

**Authors:** Alexander E. Mayer, Angel Loza-Valdes, Werner Schmitz, Jonathan Trujillo Viera, Michael Leitges, Andreas Schlosser, Grzegorz Sumara

**Affiliations:** Rudolf Virchow Center, Center for Integrative and Translational Bioimaging’ University of Würzburg, 97080 Würzburg, Germany; Nencki Institute of Experimental Biology, Polish Academy of Sciences, 3 Pasteur Street, 02-093 Warsaw, Poland; Theodor Boveri Institute, Biocenter, University of Würzburg, 97074 Würzburg, Germany; Tier 1, Canada Research Chair in Cell Signaling and Translational Medicine, Division of BioMedical Sciences / Faculty of Medicine, Craig L Dobbin Genetics Research Centre, Memorial University of Newfoundland, Health Science Centre 300 Prince Philip Drive, St. John’s, Newfoundland, Canada A1B 3V6

**Keywords:** Protein Kinase D3 (PKD3), phenylalanine hydroxylase (PAH), phenylalanine (Phe), tyrosine (Tyr), Phenylketonuria (PKU), Hepatocytes, Glucagon

## Abstract

Members of the Protein Kinase D (PKD) family (PKD1, 2, and 3) integrate hormonal and nutritional inputs to regulate complex cellular metabolism. Despite the fact that a number of functions have been annotated to particular PKDs, their molecular targets are relatively poorly explored. PKD3 promotes insulin sensitivity and suppresses lipogenesis in the liver. However, its substrates are largely unknown. Here we applied proteomic approaches to determine PKD3 targets. We identified over three-hundred putative targets of PKD3. Among them phenylalanine hydroxylase (PAH). PAH catalyses the conversion of phenylalanine to tyrosine and its activity is regulated by, phenylalanine concentration and glucagon-induced signaling. Consistently, we showed that PKD3 is activated by glucagon and promotes tyrosine levels in primary hepatocytes and liver of mice.

Taken together, our comprehensive proteomic approach established that PKD3 determine the rate of phenylalanine to tyrosine conversion in the liver. Therefore, our data indicate that PKD3 might play a role in development of diseases related to the defective tyrosine and phenylalanine metabolism.

## Introduction

Protein Kinase D (PKD) family members integrate multiple hormonal and metabolic signals to coordinate homeostasis of the organism (Kolczynska, Loza-Valdes et al., 2020, Löffler, Mayer et al., 2018, Mayer, Löffler et al., 2019, Rozengurt, 2011, Sumara, Formentini et al., 2009). The family of PKDs comprises three kinases: PKD1, PKD2, and PKD3 (Fu & Rubin, 2011, Rozengurt, 2011). PKDs share a basic structure composed of the cysteine-rich domain (CRD), essential for their affinity for their main activators phorbol esters and diacylglycerol (DAG). The pleckstrin homology domain (PH) and the c-terminal region determine the catalytic activity (Iglesias, Matthews et al., 1998, Rozengurt, Sinnett-Smith et al., 1997). PKD1 and PKD2 share the highest homology, while PKD3 kinase is the unique member of the family. PKD1 and PKD2 have been widely studied in different cellular processes such as trans-Golgi network dynamics, cell proliferation and cell migration, adipocytes function, insulin secretion as well as regulation of innate and adaptive immune cells function (Gehart, Goginashvili et al., 2012, Goginashvili, Zhang et al., 2015, Ittner, Block et al., 2012, Kolczynska et al., 2020, Löffler et al., 2018, Mayer et al., 2019, Rozengurt, 2011, Sumara et al., 2009, Zhang, Meszaros et al., 2017). The specific functions of PKD3 are still relatively unexplored. PKD3 has been implicated in tumor progression and invasiveness in breast and gastric cancers, as well as hepatocellular carcinoma (Huck, Duss et al., 2014, Yang, Xu et al., 2017, Zhang, Zhang et al., 2019). Furthermore, recent research has demonstrated that PKD3 regulates insulin sensitivity, lipid accumulation, and fibrogenesis in the liver (Mayer et al., 2019, Zhang, Liu et al., 2020). Thus, PKD3 plays a role in a wide range of cellular processes in both physiological and pathological conditions.

To date, only a few downstream targets of PKD3 have been identified. PKD3 phosphorylates G-protein-coupled receptor kinase-interacting protein 1 (GIT1) on serine 46 to regulate the localization of GIT1-paxilin complex and consequently cell shape and motility (Huck, Kemkemer et al., 2012). Moreover, ectopic expression of a constitutive active form of PKD3 (PKD3ca) in TNBC (triple-negative breast cancer cells) leads to hyperphosphorylation of S6 Kinase 1 (S6K1), a downstream target of the mechanistic target of rapamycin complex 1 (mTORC1), which is an energy sensor in the cell and sustains cell proliferation (Huck et al., 2014, Laplante & Sabatini, 2012). PKD3 also phosphorylates p65 at serine 536, a critical step for the upregulation of 6-phosphofructo-2-kinase/fructose-2,6-biphosphatase 3 (PFKFB3) and drives glycolysis in gastric cancer cells (Zhang et al., 2019). In addition, gain and loss of function studies suggest that PKD3 regulates the ERK1-MYC axis and promotes cell proliferation in cancer (Chen, Deng et al., 2008, Liu, Song et al., 2019). Finally, in hepatocytes, PKD3 suppresses insulin-dependent AKT and mTORC1/2 activation, which results in peripheral glucose intolerance and suppression of the hepatic lipid production (Mayer et al., 2019). Nevertheless, the PKD3 targets in the liver and other organs remain largely unexplored.

The liver has a major role in the regulation of glucose, lipid, and amino acids (AAs) homeostasis by regulating the adaptation to nutrient availability. In the liver, AAs are utilized to synthesize proteins and precursors for different bioactive molecules. Moreover, ammonia, a by-product of protein catabolism, is disposed as urea by the liver (Bröer & Bröer, 2017, Waterlow, 1999). Under certain physiological conditions such as fasting, the liver can utilize AAs to produce glucose or ketone bodies. This metabolic response is hormonally regulated by glucagon, which is released from the pancreatic alpha cells (Holst, Albrechtsen et al., 2017, Petersen, Vatner et al., 2017).

Phenylalanine (Phe) is an essential AA in mammals, and its conversion into tyrosine (Tyr) is crucial for the production of thyroid hormones and catecholamines. The conversion into tyrosine is tightly regulated by the enzyme phenylalanine hydroxylase (PAH), an enzyme that requires tetrahydrobiopterin (BH_4_) as a cofactor, and molecular dioxygen as a substrate (Fitzpatrick, 1999, Kaufman, 1958). Mutations in PAH lead to phenylketonuria (PKU), an abnormal accumulation of Phe in peripheral tissues (Konecki & Lichter-Konecki, 1991, Scriver, 2007, Williams, Mamotte et al., 2008). Of note, the expression of PAH is restricted to the liver and kidney, major organs involved in AAs metabolism (Hsieh & Berry, 1979). PAH activity is regulated allosterically by high intracellular levels of Phe and hormonally by glucagon and insulin. By contrast, it was recently shown that oxygen concentrations might affect PAH activity in hepatocytes due to oxygen zonation (Donlon & Kaufman, 1978b, Kaufman, 1958, Ying, Pey et al., 2010). Upon fasting, glucagon rewires liver metabolism and promotes AAs catabolism. Glucagon leads to an increase in cAMP (cyclic Adenosine Monophosphate) and activates protein kinase A (PKA), which phosphorylates PAH at serine 16 to promote its function and increase the rate of Phe to Tyr conversion (Miranda, Teigen et al., 2002).

Here we carried out a phosphoproteomic study to investigate phosphorylation events dependent on PKD3 and thus to identify putative PKD3 targets in the liver. We found over three hundred of putative substrates of PKD3, among them PAH. Consistently, in mice and primary hepatocytes overexpressing constitutive active form of PKD3 (PKD3ca), Tyr levels were elevated, while the deletion of PKD3 resulted in decreased conversion of Phe to Tyr. Moreover, we showed that glucagon signaling promotes PKD3 activation and PAH activity in hepatocytes. Finally, PKD3 is required for glucagon-induced Phe to Tyr conversion. Taken together, we have identified potential PKD3-specific substrates in hepatocytes and uncovered the function of PKD3 in the regulation of Phe and Tyr metabolism.

## RESULTS

### Unraveling putative targets of PKD3 in hepatocytes using substrate motif-specific antibodies

Previous research has delineated the role of PKD3 in the regulation of hepatic glucose and lipid metabolism (Mayer et al., 2019). However, the phosphorylation targets of PKD3 in the liver remain elusive. To unravel the putative targets of PKD3 in the liver we have utilized primary hepatocytes derived from PKD3 knockout mice and transduced these cells with adenovirus to overexpress either EGFP or PKD3ca (Fig. 1A). Subsequently, protein lysates were isolated, and used for pull-down assays. For this, we utilized PKD substrate motif specific antibodies. PKD kinases recognize the consensus AAs motif sequence LxRxx[S*/T*] (where L – leucine, R – arginine, S – serine, T – threonine, and x – any AA) within their putative targets (Döppler, Storz et al., 2005, Franz-Wachtel, Eisler et al., 2012). Importantly, the arginine (R) in position −3 in relation to the phospho-acceptor is essential, whereas leucine (L) in −5 position might be in some cases replaced by other amino acids e.g. valine (V) or isoleucine (I) (Döppler et al., 2005). At first, phospho-(Ser/Thr) PKD substrate LxRxx[S*/T*] antibody was used for immunoprecipitation to enrich proteins that have a phosphorylated PKD motif in lysates from primary hepatocytes deficient for PKD3 expressing either EGFP control or PKD3ca (Fig. 1A). Overexpression of PKD3ca showed an increase in proteins with a PKD motif (Fig. 1B). Proteins with phosphorylated PKD motif were immunoprecipitated and characterized by mass spectrometry. 84 proteins were significantly enriched (significance of 1 or 2) in lysates from PKD3ca hepatocytes compared to the EGFP ones (Fig. 1C, D and Supplementary table 1). The protein with highest enrichment induced by PKD3ca was PKD3 itself, suggesting that PKD3 might be subjected to auto-phosphorylation. Although leucine in −5 position is frequently present in PKD motifs, other targets also have valine (V) or isoleucine (I) in −5 position (Döppler et al., 2005). Therefore, to complement our approach, we performed a similar experiment utilizing an antibody only partially specific for PKD motif Rxx[S*/T*]. Overexpression of PKD3ca led to an increase in proteins that have a phosphorylated Rxx[S*/T*] motif (Fig. 2A). Subsequently, proteins were immunoprecipitated, identified and quantified by mass spectrometry. 226 proteins were significantly enriched (significance of 1 or 2) in lysates from PKD3ca hepatocytes compared to the EGFP expressing ones (Fig. 2B, C and Supplementary table 3). Of note, using antibody against motif Rxx[S*/T*] we identified almost three times more putative targets of PKD3 compared to antibody against LxRxx[S*/T*] motif, suggesting that PKD3 can frequently phosphorylate imperfect consensus site.

**Figure 1:**
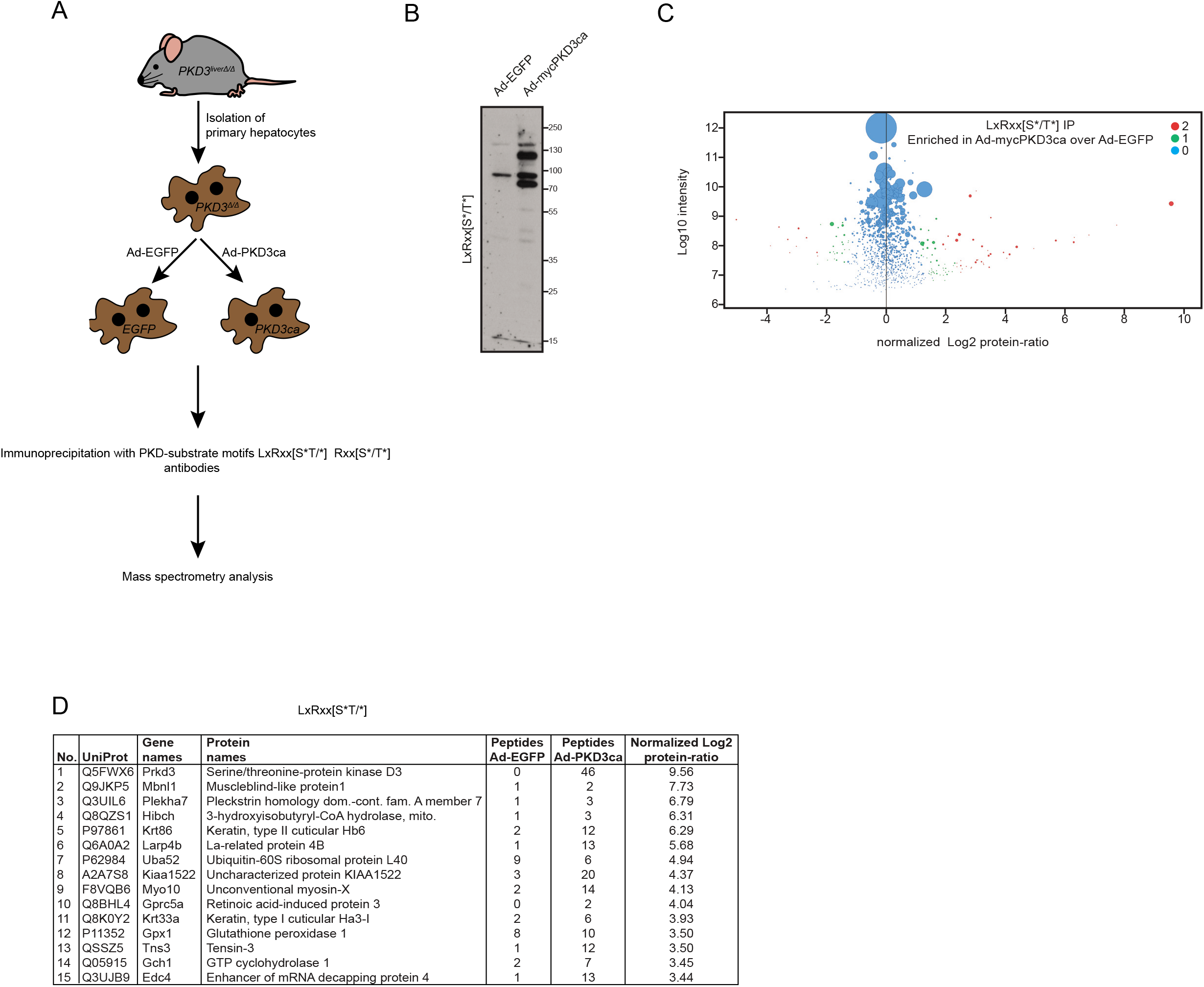
Identification of PKD3 substrates by immunoprecipitation using PKD substrate motif antibody LxRxx[S*/T*]. (A) Experimental design. Primary hepatocytes isolated from PKD3-deficient mice were transduced by adenovirus containing EGFP (controls) or the constitutive active form of PKD3 (PKD3ca). IP with PKD-substrate motif antibodies was performed on cell extracts and followed by mass spectrometry analysis. (B) WB analysis of protein lysates from PKD3-deficient primary hepatocytes transduced with either adenovirus expressing control EGFP (Ad-EGFP) or constitutive active PKD3 (Ad-mycPKD3ca) using PKD-substrate motif LxRxx[S*/T*]-specific antibody (n=3 independent experiments). (C) Scatter plot of the statistical significance of log2 transformed protein ratios versus log10-transformed LFQ intensities between control and PKD3ca expressing hepatocytes. Enriched proteins are indicated by red (2, significantly enriched) or green (1, potentially enriched) dots, blue ones are not enriched (0, not enriched). (D) 15 most enriched proteins identified by mass spectrometry showing number, UniProt accession number, gene names, protein names, peptide count in EGFP and PKD3ca samples, respectively, and log2 transformed LFQ protein ratio are shown in the table.

**Figure 2:**
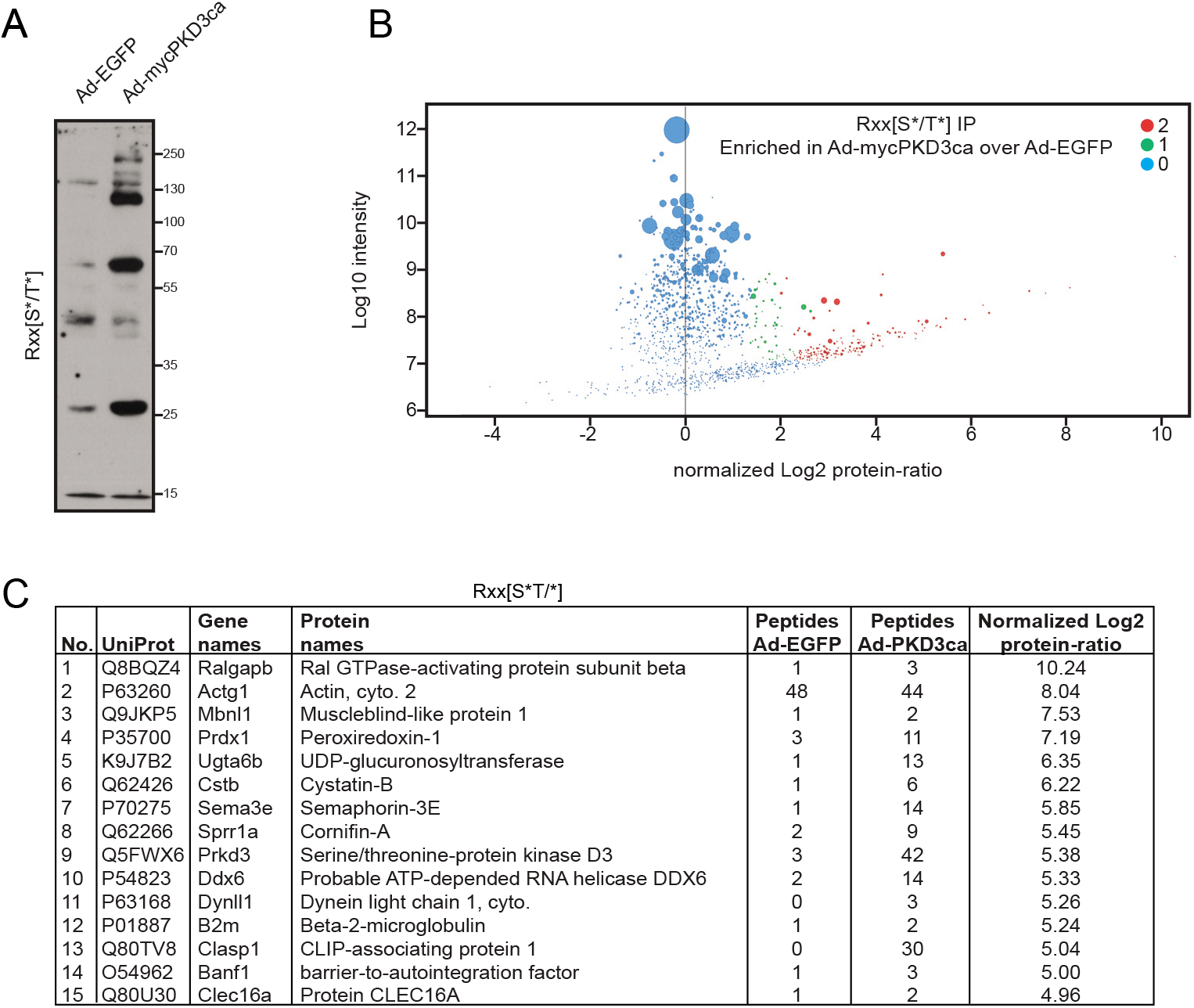
Identification of PKD3 targets in hepatocytes by IP with Rxx[S*/T*] antibody. (A) WB using Rxx[S*/T*] motif antibody on lysates from PKD3-deficient hepatocytes expressing control EGFP (Ad-EGFP) or constitutive active form of PKD3 (Ad-mycPKD3ca). (B) Scatter plot of the statistical significance of log2 transformed protein ratios versus log10-transformed LFQ intensities between control and PKD3ca expressing hepatocytes. Enriched proteins are indicated by red (2, significantly enriched) or green (1, potentially enriched) dots, blue ones are not enriched (0, not enriched). (C) 15 most enriched proteins identified by mass spectrometry in the experiment from C, showing number, UniProt accession number, gene names, protein names, peptide count in EGFP and PKD3ca samples, respectively, and log2 transformed LFQ protein ratio.

### Comparative analysis of putative PKD3 targets identified using antibodies against LxRxx[S*/T*] and Rxx[S*/T*] motifs

Our mass spectrometry screening identified 84 and 226 proteins significantly enriched using antibodies against LxRxx[S*/T*] and Rxx[S*/T*], respectively (Fig. 3A). Of note, 24 proteins were enriched in both screenings (Fig. 3A and C). In silico analysis revealed that 55% of proteins identified using antibody against LxRxx[S*/T*] motif, have at least one putative PKD consensus side resembling the sequence [L/V/I]xRxx[S*/T*]. Similarly, also 55% of proteins enriched from hepatocytes expressing PKD3ca using antibody against Rxx[S*/T*] had at least one [L/V/I]xRxx[S*/T*] motif in their sequence (Fig. 3B). 12 of the proteins which had in their sequence a [L/V/I]xRxx[S*/T*] motif, were enriched using both antibodies, against Rxx[S*/T*] and LxRxx[S*/T*] (Fig. 3B). In silico analysis also revealed that these 12 proteins have in total 30 putative PKD motifs. Further analysis of the 30 putative PKD motifs carried out in phosphosite.org repository showed that 11 sites of the motifs were previously reported. Noteworthy, 3 of the motifs were identified in the field of PKDs (also PKA signaling), namely LsRklS16 for phenylalanine hydroxylase (PAH), and LtRqkS3894 as well as LtRqlS5407 for dystonin (DST) (Fig. 3D). Moreover, a prediction tool for biological processes (ARCHS4) suggests that PKD3 signaling might influence catabolic and metabolic activity of PAH (Fig. 3E). PAH converts Phe to Tyr (Kaufman, 1958), therefore these data suggest that PKD3 might also regulate AAs metabolism in liver.

**Figure 3:**
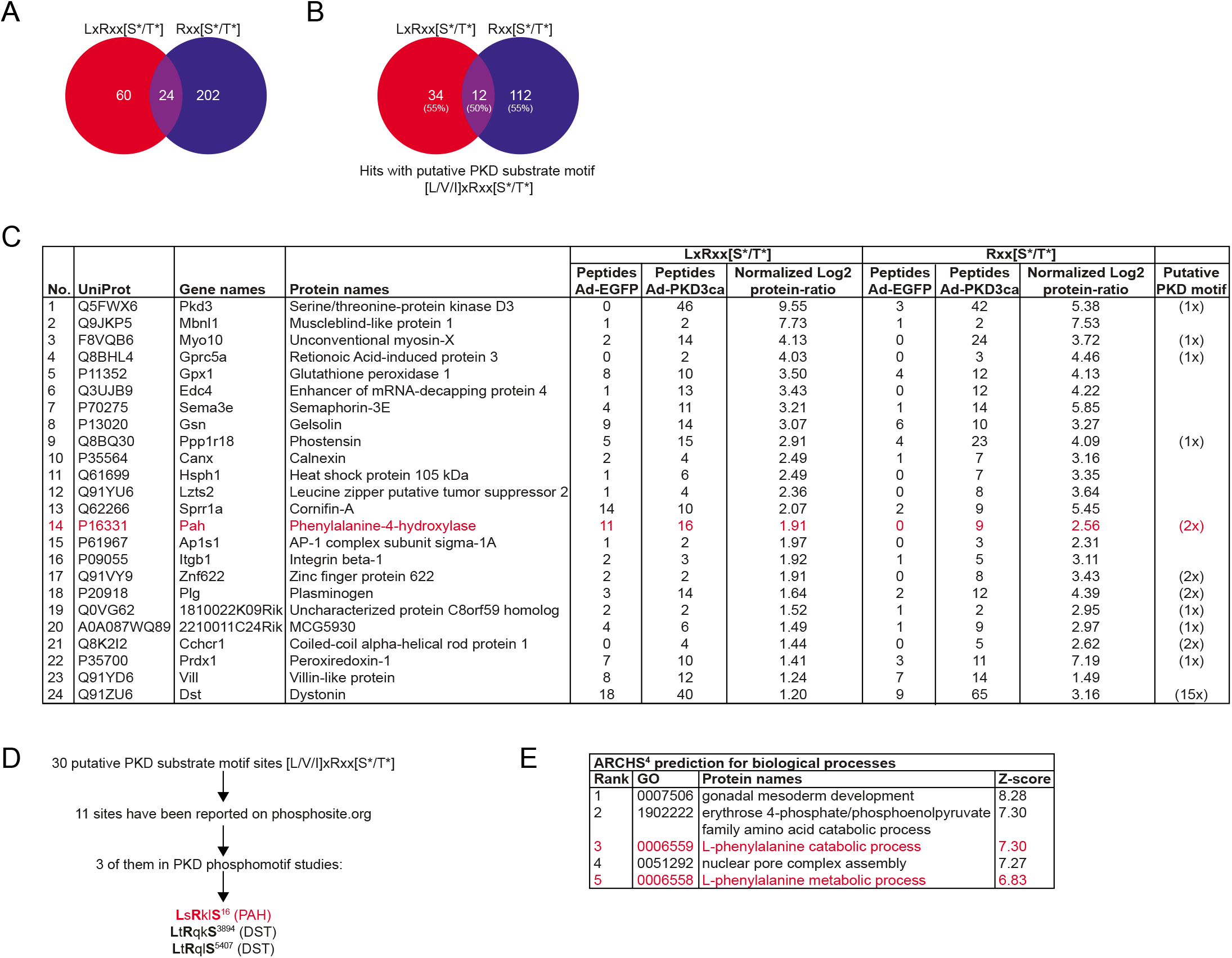
Comparative analysis of putative PKD3 targets identified using different motif antibodies. (A) Venn diagram of common putative substrates that were significantly enriched by both antibodies LxRxx[S*/T*] and Rxx[S*/T*]. (B) Computational identification of putative PKD motifs [L/V/I]xRxx[S*/T*] among putative substrates identified by both antibodies (percentage of proteins possessing a putative motif in brackets). Results were obtained using ExPASy ScanProsite tool. (C) List of 24 proteins enriched by both antibodies, LxRxx[S*/T*] and Rxx[S*/T*] with UniProt accession numbers, gene and protein names, peptide count and log2 transformed LFQ protein ratio for LxRxx[S*/T*] and Rxx[S*/T*] immunoprecipitations, respectively, and the amount of putative PKD motifs for each protein. (D) Flow of computational identification of putative PKD substrates by comparing the PKD motifs within proteins using phosphosite.org repository. (E) Prediction of biological processes potentially regulated by PKD3 using ARCHS^4^.

### PKD3 signaling determines tyrosine levels in liver

To test whether PKD3 regulates PAH phosphorylation in hepatocytes, we transduced primary hepatocytes with increasing amounts of adenoviruses expressing either EGFP or PKD3ca. Subsequently, protein lysates were used to evaluate PAH migration on the SDS PAGE followed by Western blotting. Interestingly, overexpression of PKDca leads to an upshift of PAH signal and appearance of the second band, which is specific for this protein. Interestingly, the upshift of PAH was more pronounced when hepatocytes were transduced with an increasing amount of PKD3ca (Fig. 4A). Furthermore, to explore the physiological role of PKD3 in Phe metabolism, we cultivated hepatocytes expressing either EGFP or PKD3ca in the medium deprived of Phe and Tyr. Following Phe/Tyr starvation, we supplemented the cell culture medium of hepatocytes with increasing amounts of Phe and determined the tyrosine levels in the cells. In the cells, which were not supplemented with Phe, expression of PKD3ca resulted in the most pronounced increase in Tyr levels compared to control hepatocytes. Supplementation of Phe in the medium resulted in increased concentrations of Tyr levels in control hepatocytes. However, at each of the tested conditions, Tyr levels were significantly higher in the cells expressing PKD3ca but not increasing further upon addition of Phe (Fig. 4B). This suggests that PKD3 promotes the conversion of Phe to Tyr in hepatocytes. To test whether PKD3 regulates levels of Tyr in the complex in vivo situation, we measured Tyr levels in mice expressing PKD3ca specifically in hepatocytes (Mayer et al., 2019). Of note, mice overexpressing PKD3ca presented higher levels of Tyr in hepatic extracts than corresponding control animals (Fig. 4C). Moreover, as revealed by metabolomics analysis, mice overexpressing PKD3ca presented also higher Tyr to Phe ratio compared to control animals, while the levels of other AAs were not altered (Fig. 4D and Supplementary table 2). Altogether, these results suggest that PKD3 has a key role in the conversion of Phe into Tyr in the liver.

**Figure 4:**
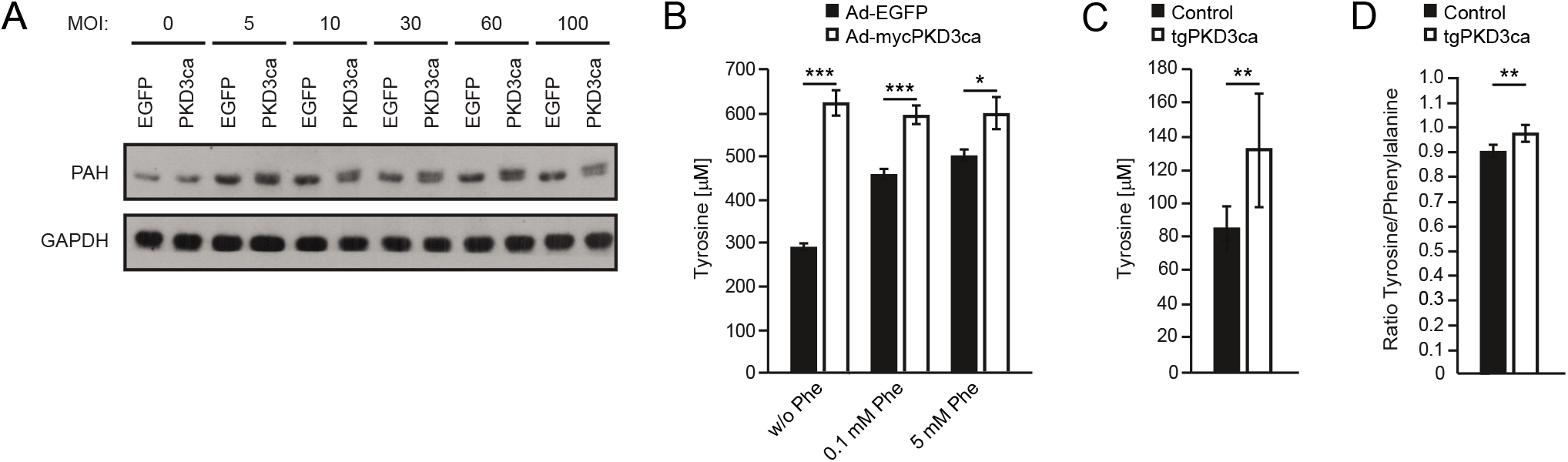
PKD3 signaling controls PAH activity and determines cellular levels of Tyr. (A) PAH expression and shifting analyzed in hepatocytes transduced with increasing amounts of adenovirus expressing EGFP or PKD3ca at indicated MOIs using WB. (B) Phe to Tyr conversion assay in primary hepatocytes expressing either EGFP or PKD3ca. Cells were depleted from Phe and Tyr in the medium for 1 h before stimulation and incubated with 0, 0.1, or 5 mM Phe for 1 h. (C) Tyr levels in livers from control mice and mice overexpressing PKD3ca (n=12 and n=16). (D) Tyr to Phe ratio in liver tissues from control mice and mice overexpressing PKD3ca (n=12 and n=16).

### A glucagon-PKD3 axis determines amino acid metabolism in the liver

Seminal papers in the early 1970s demonstrated that hepatic PAH activity is hormonally regulated by glucagon via PKA signaling (Abita, Milstien et al., 1976a, Donlon & Kaufman, 1978b). PKA phosphorylates Ser16 of PAH upon glucagon stimulation, which increases the affinity of PAH for its main substrate, the amino acid Phe (Miranda et al., 2002). In addition, glucagon promotes formation of DAG, a well-known activator of PKD3 (Hermsdorf, Dettmer et al., 1989, L Rodgers, 2012). Glucagon is a classical hormone induced under fasting conditions that rewires metabolism in the liver primarily via PKA (Pilkis, Claus et al., 1988, Pilkis & Granner, 1992). Thus, in order to investigate whether glucagon also governs PKD activity and AAs metabolism, we starved primary hepatocytes from all AA for one hour, then cultured them in media containing either no amino acids (w/o AA), all amino acids (AA), all amino acids except Phe and Tyr (w/o Phe, Tyr), Phe exclusively, or Phe and Tyr exclusively for additional hour and stimulated them with glucagon for 5 or 20 minutes. Glucagon promotes phosphorylation of PKD3-Ser731/735 independent of amino acid stimulation. In addition, stimulation of primary hepatocytes with all AAs or with all AA except Phe and Tyr was sufficient to activate S6K-Thr389 as well as 4E-BP1-Thr37/46 and Ser65, which was reduced upon glucagon stimulation in a time-dependent manner after 5 and 20 min (Fig. 5A). Importantly, similarly to cells expressing PKD3ca, stimulation of hepatocytes with glucagon resulted in an upshift of PAH on the Western blot, probably resembling increased phosphorylation of this enzyme (Fig. 5A). All of these suggest that PKD3 might be required for glucagon-induced conversion of Phe to Tyr. Indeed, stimulation of primary control hepatocytes with glucagon resulted in increased Tyr levels, but in the cells-derived from PKD3-deficient mice, glucagon failed to increase Tyr levels (Fig. 5B). All of these data suggest that activation of PKD3 by glucagon is required for induction of Phe to Tyr conversion.

**Figure 5:**
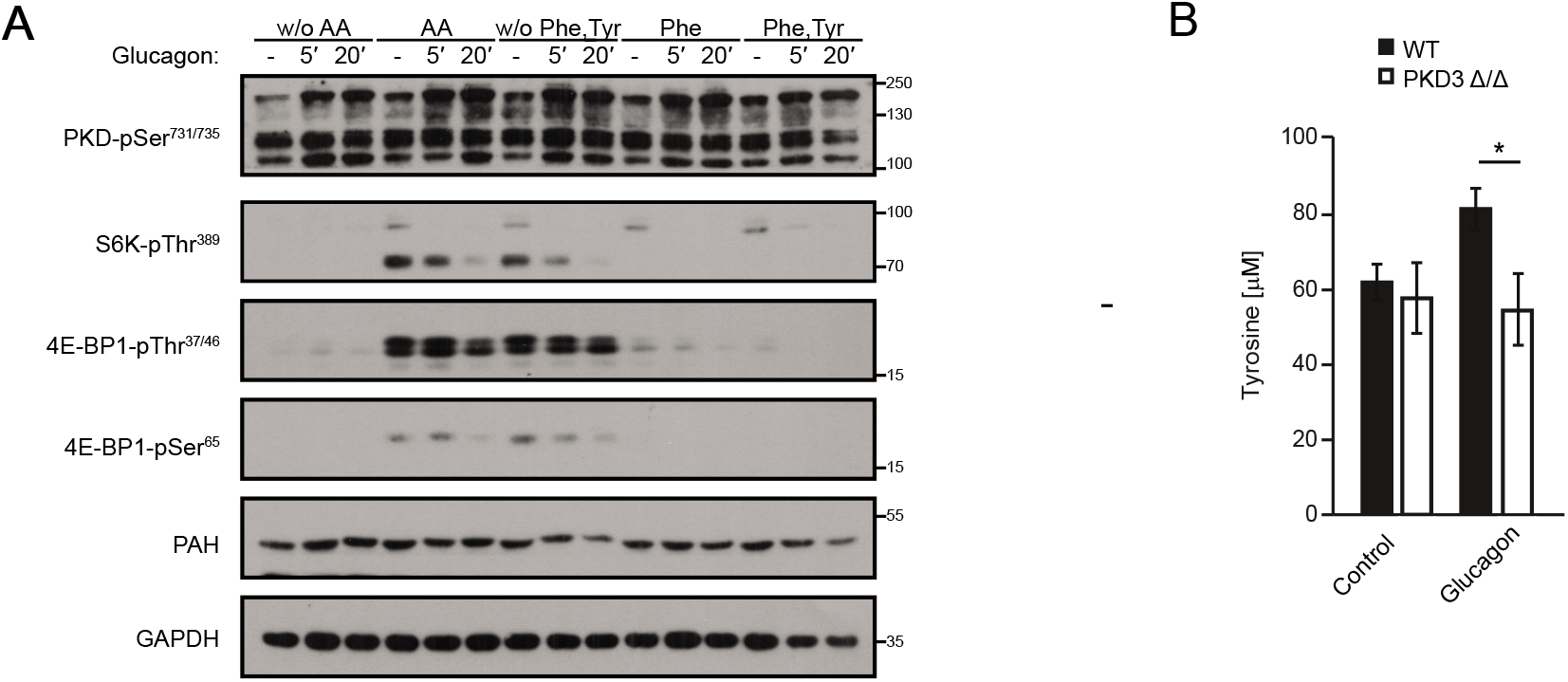
A glucagon-PKD3 axis determines amino acids metabolism in the liver. (A) WB analysis of indicated proteins in primary hepatocytes deprived from all AA for one hour, then cultured in media containing either no amino acids (w/o AA), all amino acids (AA), all amino acids except Phe and Tyr (w/o Phe, Tyr), Phe exclusively, or Phe and Tyr exclusively for additional hour and stimulated with glucagon for 5 or 20 minutes. (B) Intracellular phenylalanine to tyrosine conversion assay in primary hepatocytes isolated from control or PKD3-deficient mice and stimulated with glucagon for 20 minutes. Data are presented as mean ± SEM. *P>0.05, ***P>0.001 (one way ANOVA with post hoc Tukey’s test).

## Discussion

Recent studies established PKD3 function in the regulation of hepatic lipid and glucose metabolism (Mayer et al., 2019). However, the direct targets of PKD3 in hepatic cells remained elusive. Utilizing proteomic approach on primary hepatocytes deficient for PKD3 and re-expressing PKD3ca we identified over three-hundred putative targets of PKD3. Importantly, this approach resulted in the identification of the novel function of PKD3 in the regulation of hepatic metabolism of AAs. Namely, we showed that PKD3 promotes the conversion of Phe to Tyr by activating PAH, a rate-limiting enzyme in this process.

In our studies, we utilized two complementary proteomic approaches. For pull-downs, we used two antibodies: against the full phospho-motif sequence of PKD LxRxx[S*/T*] and antibody against part of the PKD phospho-motif sequence Rxx[S*/T*]. Importantly, pulldowns using an antibody against part of the PKD motif revealed more of the potential targets of PKD3 than antibody against full sequence targeted by PKDs. This indicates that in a large number of PKD3 target proteins AA at the position −5 in relation to the phospho-acceptor AA might be other than leucine. Whether we used for pull-downs antibody against LxRxx[S*/T*] or Rxx[S*/T*] motif, almost 50% of identified proteins did not present in their sequence the consensus motif for PKDs. These might indicate that in our pull-downs we have also fished out proteins, which are interacting with the targets of PKD3 but are not phosphorylated by PKD3 themselves.

In total, 12 proteins were identified by both pull-downs and possess one or more PKD phosphorylation motifs in their sequence. Among them, we found PAH as a target for PKD3, which we confirmed by classical WB. In line with our findings, a computational analysis to predict gene functions suggests that PKD3 might be involved in phenylalanine metabolism. PAH has two putative PKD motifs (LsRklS16 and IpRpfS411). It was shown that Ser16 is phosphorylated upon glucagon stimulation via PKA activation (Abita, Milstien et al., 1976b, Donlon & Kaufman, 1978a). Furthermore, the classical PKA motif is RRx[S*/T*] and has in - 2 position of Ser16 a lysine (K) and not arginine (R), which can also serve as a PKD motif (Pinna & Ruzzene, 1996). Therefore, it is plausible that glucagon also might affect hepatic PAH activity via PKD3 signaling leading to changes in Tyr concentration. Phosphorylation of PAH on Ser16 increases the affinity of this enzyme to Phe (Arun, Kaddour-Djebbar et al., 2011, Doskeland, Martinez et al., 1996). Consistently, primary hepatocytes overexpressing PKD3ca presented higher levels of Tyr especially when cells were starved from Phe. Importantly, transgenic mice overexpressing PKD3ca had higher levels of Tyr in the liver as well as a higher Tyr to Phe ratio than littermate controls.

Canonical activation of PKA in the liver promotes amino acid catabolism (Holst et al., 2017). However, our findings suggest that glucagon also promotes the activation of PKD3 in primary hepatocytes independently of AA stimulation. Moreover, activation of PKD3 is required for the induction of conversion of Phe to Tyr by glucagon. All of these findings suggest that glucagon might act in liver also in a PKD3-dependent manner. Interestingly, our experiments provide a possible link between glucagon signaling and mTORC1 pathway since glucagon stimulation reduces drastically the activity of S6K-Thr389 as well as 4E-BP1-Thr37/46 and Ser65, which are crucial components of mTORC1.

As mentioned above, PKD3 regulates insulin signaling and lipogenesis in the liver by modulation of mTORC1, mTORC2, and AKT signaling (Mayer et al., 2019). Our current proteomic approach identified several potential targets of PKD3, which could explain the suppression of mTORC1, mTORC2, and AKT signaling by PKD3. For instance, Ral GTPase activating protein non-catalytic beta subunit (RalGAPβ), which can control mTORC1 activity in response to insulin stimulation (Chen, Leto et al., 2011, Martin, Chen et al., 2014) was the most enriched protein in the Rxx[S*/T*]-motif antibody-based pull-down. Moreover, Tuberous sclerosis (TSC) 1 and 2, which also control mTORC1 and mTORC2 activity (Ben-Sahra & Manning, 2017), were also found using our strategy to be a putative target of PKD3. Interestingly, the distal downstream target of mTORC2, NDRG1, was also found to be a putative target of PKD3. Since previous studies showed that NDRG1 promotes lipogenesis (Cai, El-Merahbi et al., 2017), this might also explain PKD3-dependent lipogenesis in the liver. However, these putative targets of PKD3 require confirmation and the detailed functions of PKD3-dependent phosphorylation needs to be identified.

In different cancer cell types, GIT1, S6K1 and PFKFB3 have been identified as targets of PKD3 (Huck et al., 2014, Huck et al., 2012, Laplante & Sabatini, 2012, Zhang et al., 2019). However, these proteins did not appear in our pull-downs. This might indicate that in hepatocytes PKD3 phosphorylates different set of the proteins that in respective cancer cell types.

In summary, in our study we identified a plethora of putative targets for PKD3 in the liver. Among them PAH, which suggests that PKD3 plays a role in AAs metabolism. We confirmed that PKD3 promotes the conversion of Phe to Tyr in response to the glucagon stimulation. Moreover, we have identified numerous putative targets, which might suggest the role of PKD3 in the regulation of lipid metabolism or in insulin-dependent signaling.

## Materials and Methods

### Primary hepatocyte isolation, culture and infection

Primary mouse hepatocytes were prepared as described previously (Mayer et al., 2019). All relevant mouse models of PKD3-deficiency or overexpression were also described in (Mayer et al., 2019). All animal studies were approved by the local animal welfare authorities (Regierung von Unterfranken) with the animal protocol no. AK55.2-2531.01-124/13. All mouse primary hepatocytes were infected 4 to 6 hours after plating with adenoviruses expressing either enhanced green fluorescent protein (EGFP) (Ad-EGFP) or a constitutively active form of PKD3 (Ad-mycPKD3ca) at a multiplicity of infection (MOI) of 10. Medium was replaced the following morning, and cells were used for experiments 36 to 48 hours after infection. Transduction efficiency, which was 100%, was assessed by analyzing the expression of the EGFP reporter (which was present in all adenoviruses).

### Phenylalanine conversion assay

Primary hepatocytes were fasted in DMEM without phenylalanine (Phe) and tyrosine (Tyr) for 1 h. Afterwards, cells were stimulated with either 0, 0.1 or 5 mM Phe for 1 h (500 μL/well, 12-well plate). Next, the cells were lysed in 120 μL lysis buffer followed by centrifugation at 10000*g* for 10 min at 4°C. Tyrosine was quantified using the Tyrosine Assay Kit (ABNOVA) according to the manufacturer’s protocol.

### Amino acids and glucagon stimulation

Primary hepatocytes were fasted in DMEM without AAs for 1 h. Then they were cultured with either no amino acids, with all amino acids, all amino acids except Phe and Tyr, with Phe exclusively, or Phe and Tyr exclusively for 1 h. DMEM without AA and DMEM without Phe and Tyr were supplemented with respective amounts of glucose, serine, glycine, Phe, and Tyr. Hepatocytes were stimulated with glucagon for 0, 5, and 20 min.

### Immunoprecipitation (IP) and Western blot (WB) analysis

IP was performed on hepatocytes lysates using antibodies against LxRxx[S*/T*] and Rxx[S*/T*] phospho-motifs (both Cell Signaling) with Pierce Protein A/G Magnetic Beads according to the manufacturer’s protocol. Briefly, 4 mg of protein (2 mg/mL) and 30 μg of antibody were used for each IP. Samples were eluted in 1x NuPAGE lithium dodecyl sulfate (LDS) sample buffer supplemented 60 mM DDT for 10 min at 95°C. Beads were magnetically separated from the immunoprecipitated product, which was further analyzed on WB or by Mass spectrometry.

### Mass Spectrometry analysis

Gel electrophoresis and in-gel digestion were carried out according to the standard procedures.

An Orbitrap Fusion equipped with a PicoView ion source and coupled to an EASY-nLC 1000 were used for NanoLC-MS/MS analyzes.

MS and MS/MS scans were both obtained using an Orbitrap analyzer. The raw data was processed, analyzed, and quantified using the MaxQuant software. *Label-free quantification* (LFQ) intensities were used for protein quantification. Proteins with less than two identified razor and unique peptides were excluded. Data imputation was performed with values from a standard normal distribution with a mean of the 5% quantile of the combined log10-transformed LFQ intensities and a standard deviation of 0.1. Log2 transformed protein ratios of sample versus control with values outside a 1.5x (potential, significance 1) or 3x (extreme, significance 2) *interquartile range* (IQR), respectively, were considered as significantly enriched.

### Metabolomics analysis (HPLC)

For AAs analysis, pieces of mouse liver were homogenized in 69 vol. methanol/H2O (80/20, v/v) containing 3.5 μM lamivudine as external standard using an ultraturrax. The resulting homogenate was centrifuged (2 min. max rpm) and 600 μl supernatant was applied to an activated and equilibrated RP18 SPE-column (activation with 1 ml acetonitrile and equilibration with 1 ml methanol/H_2_O (80/20, v/v) (Phenomenex Strata C18-E, 55 μm, 50 mg/1 ml, Phenomenex Aschaffenburg, Germany). The resulting eluate was evaporated at room temperature using a vacuum concentrator. The evaporated samples were re-dissolved in 100 μl of 5 mM NH4OAc in acetonitrile/ H2O (25/75, v/v). The equipment used for LC/MS analysis was a Thermo Scientific Dionex Ultimate 3000 UHPLC system hyphenated with a Q Exactive mass spectrometer (QE-MS) equipped with a HESI probe (Thermo Scientific, Bremen, Germany). LC parameters: Mobile phase A consisted of 5 mM NH4OAc in acetonitrile/H_2_O (5/95, v/v), and mobile phase B consisted of 5 mM NH4OAc in acetonitrile/H_2_O (95/5, v/v). Chromatographic separation of AAs was achieved by applying 3 μl of dissolved sample on a SeQuant ZIC-HILIC column (3.5 μm particles, 100 × 2.1 mm) (Merck, Darmstadt, Germany), combined with a Javelin particle filter (Thermo Scientific, Bremen, Germany) and a SeQuant ZIC-HILIC precolumn (5 μm particles, 20 × 2 mm) (Merck, Darmstadt, Germany) using a linear gradient of mobile phase A (5 mM NH4OAc in acetonitrile/H_2_O (5/95, v/v)) and mobile phase B (5 mM NH4OAc in acetonitrile/H_2_O (95/5, v/v)). The LC gradient program was 100% solvent B for 2 min, followed by a linear decrease to 40% solvent B within 16 min, then maintaining 40% B for 6 min, then returning to 100% B in 1 min and 5 min 100% solvent B for column equilibration before each injection. The column temperature was set to 30 °C, the flow rate was maintained at 200 μL/min. The eluent was directed to the ESI source of the QE-MS from 1.85 min to 20.0 min after sample injection. MS Scan Parameters: Scan Type: Full MS, Scan Range: 69.0 - 1000 m/z, Resolution: 70,000, Polarity: Positive and Negative, alternating, AGC-Target: 3E6, Maximum Injection Time: 200 ms HESI Source Parameters: Sheath gas: 30, auxiliary gas: 10, sweep gas: 3, spray voltage: 2.5 kV in pos. mode and 3.6 kV in neg. mode, Capillary temperature: 320 °C, S-lens RF level: 55.0, Aux Gas Heater temperature: 120 °C. Data Evaluation: Peaks corresponding to the calculated amino acid masses (MIM +/− H+ ± 2 mMU) were integrated using TraceFinder software (Thermo Scientific, Bremen, Germany). Alternatively, (for Fig. 4B and C and Fig. 5B) commercially available kit was used for Tyr quantification (Cell Biolabs - Biocat).

## ACKNOWLEDGMENTS

We thank Dr. Olga Sumara for critical reading of our manuscript. This study was funded by European Research Council (ERC) Starting Grant SicMetabol (no. 678119), Emmy Noether grant Su 820/1-1 from the German Research Foundation, EMBO Installation Grant from European Molecular Biology Organization (EMBO), and the Dioscuri Centre of Scientific Excellence. The program was initiated by the Max Planck Society (MPG), managed jointly with the National Science Centre and mutually funded by the Ministry of Science and Higher Education (MNiSW) and the German Federal Ministry of Education and Research (BMBF).

---

**Supplementary Table 1.**
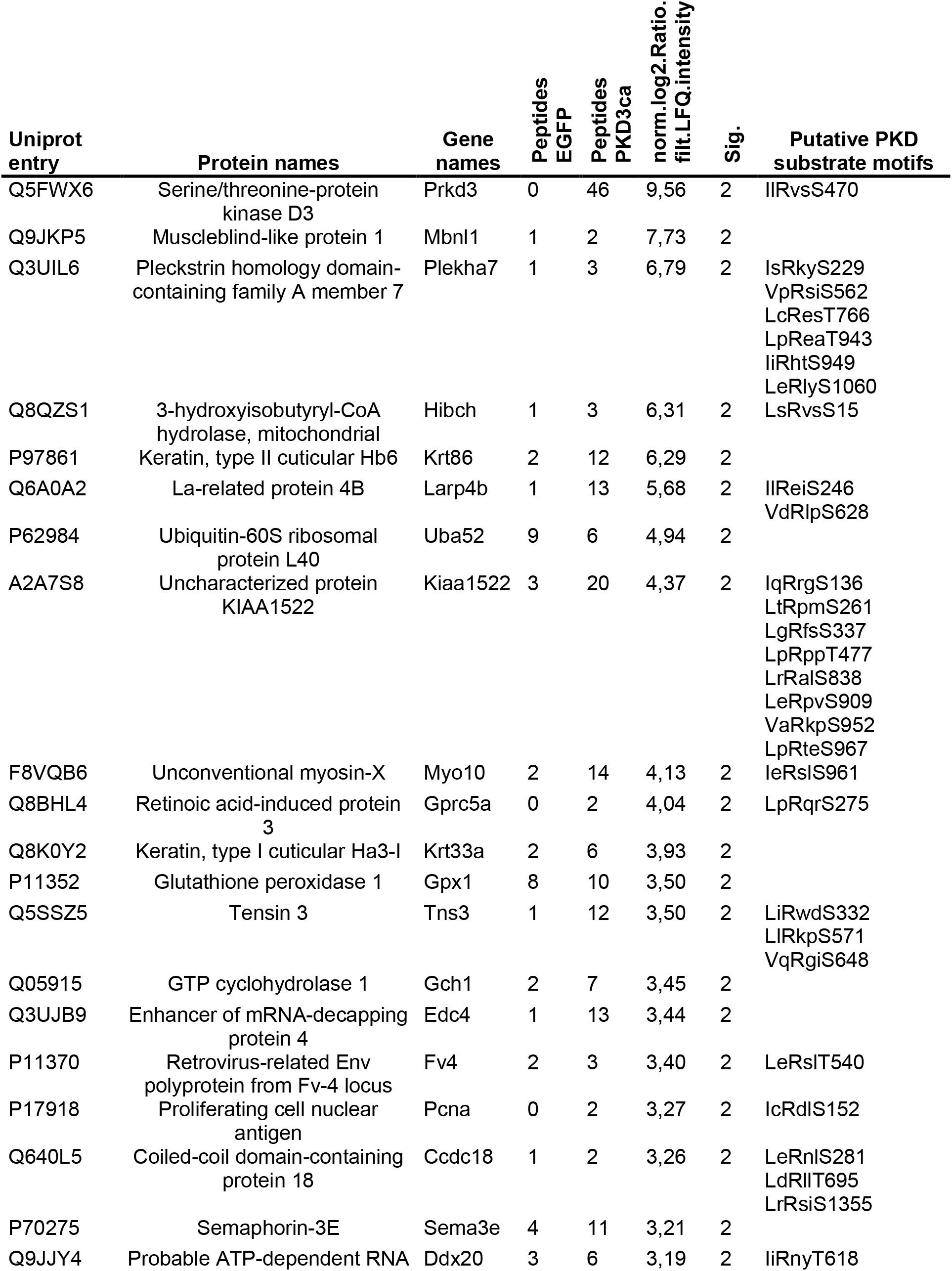
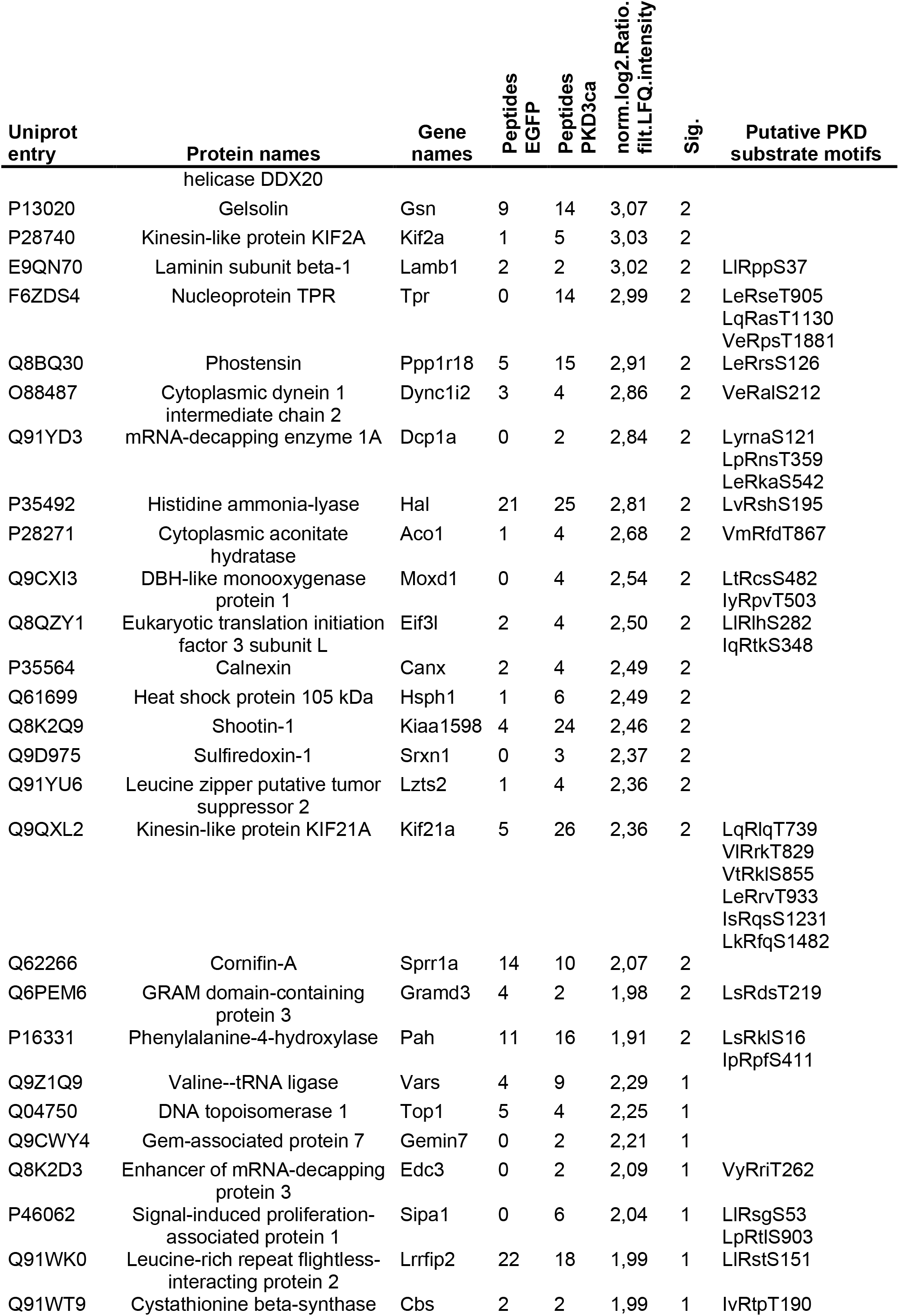
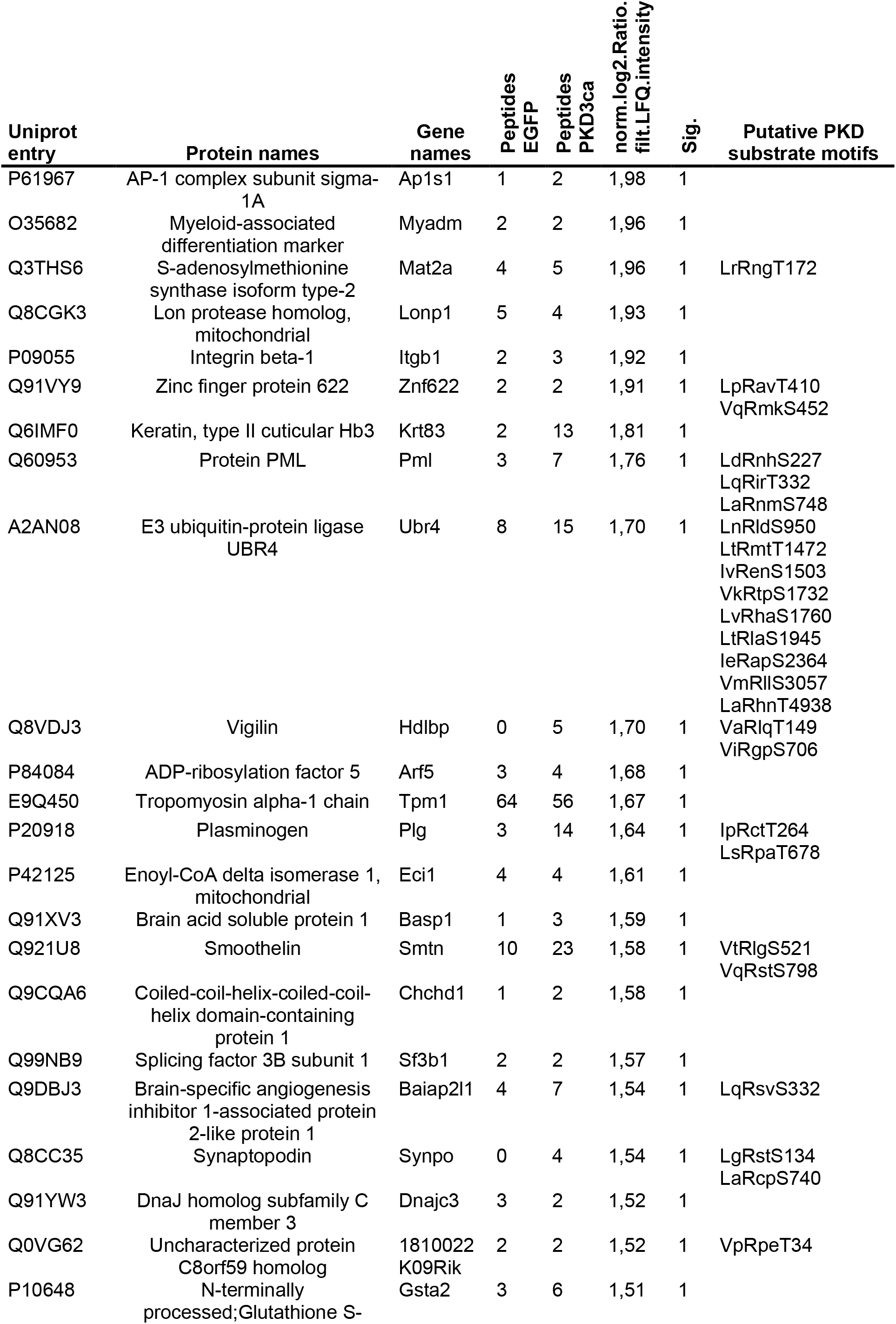
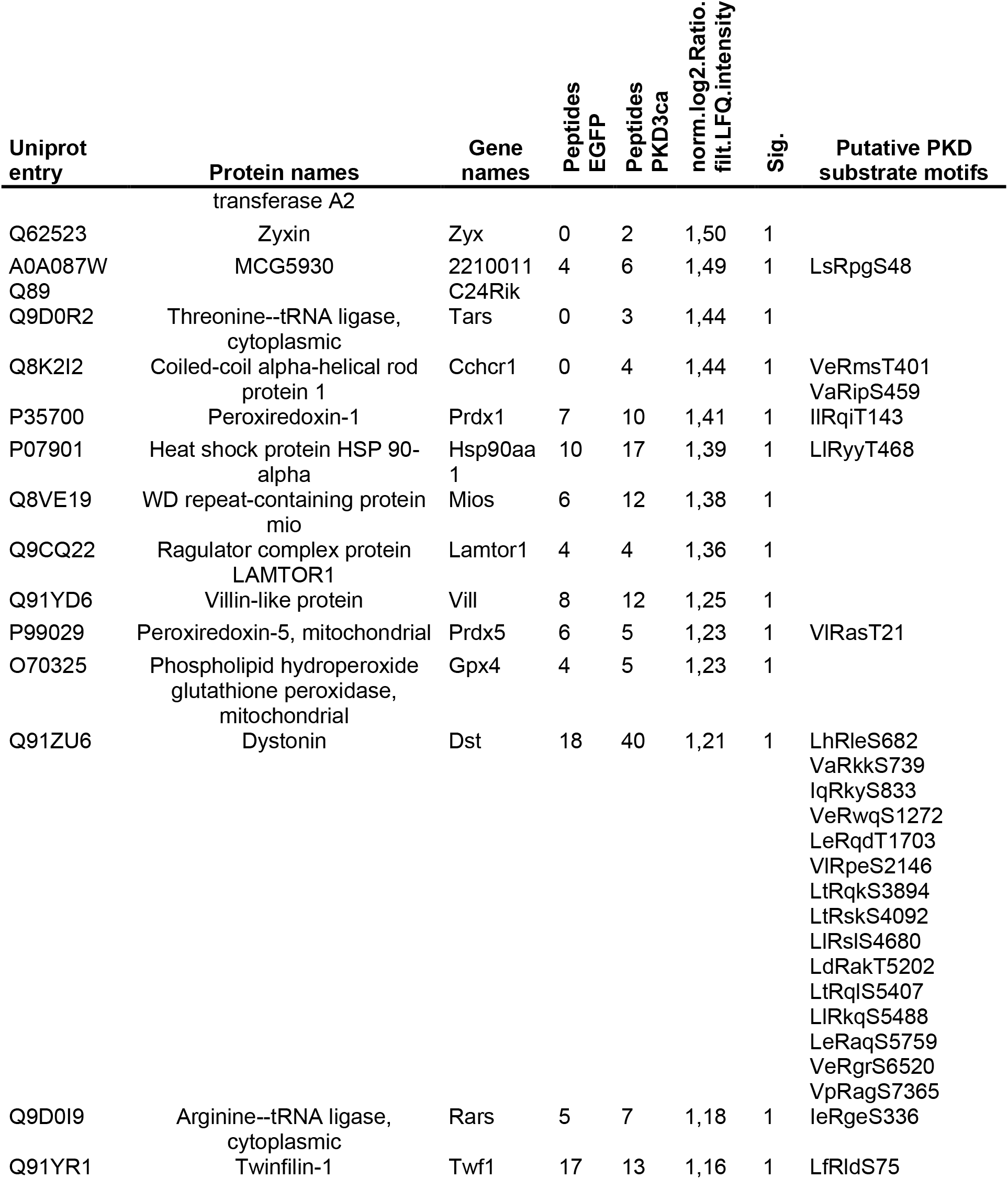
Proteins identified by MS from IP with LxRxx[S*/T*] antibody

**Supplementary Table 2.**
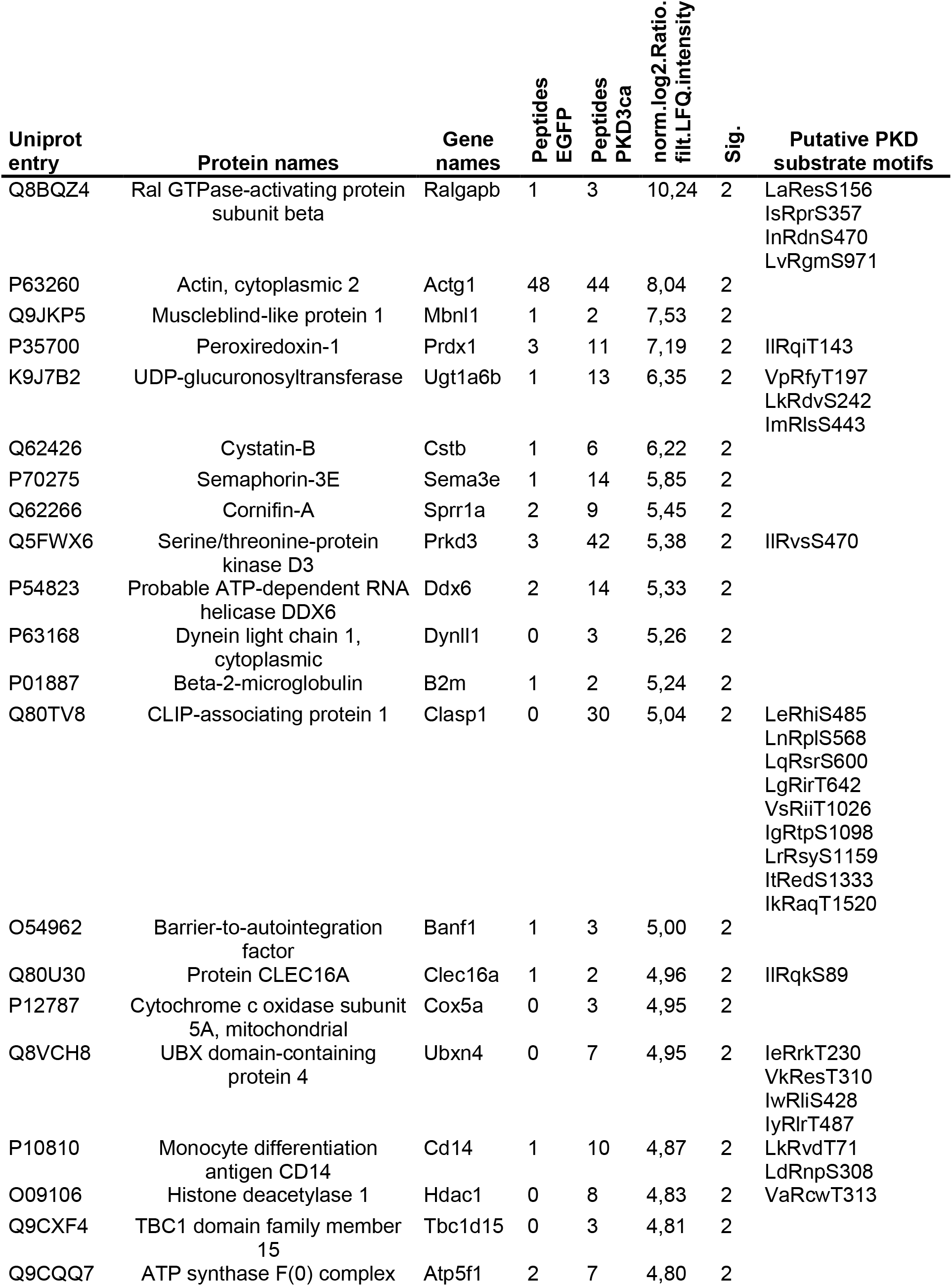
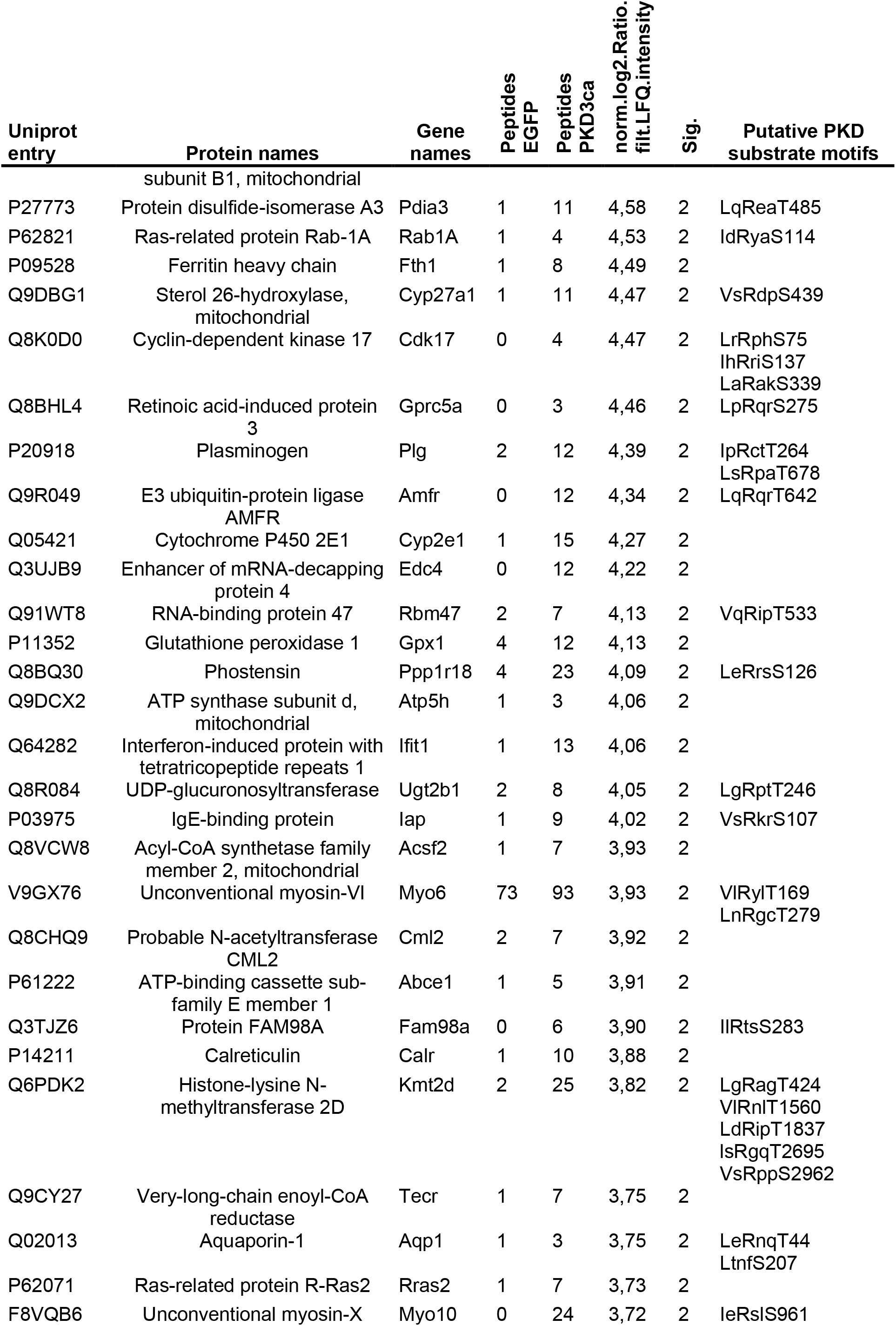
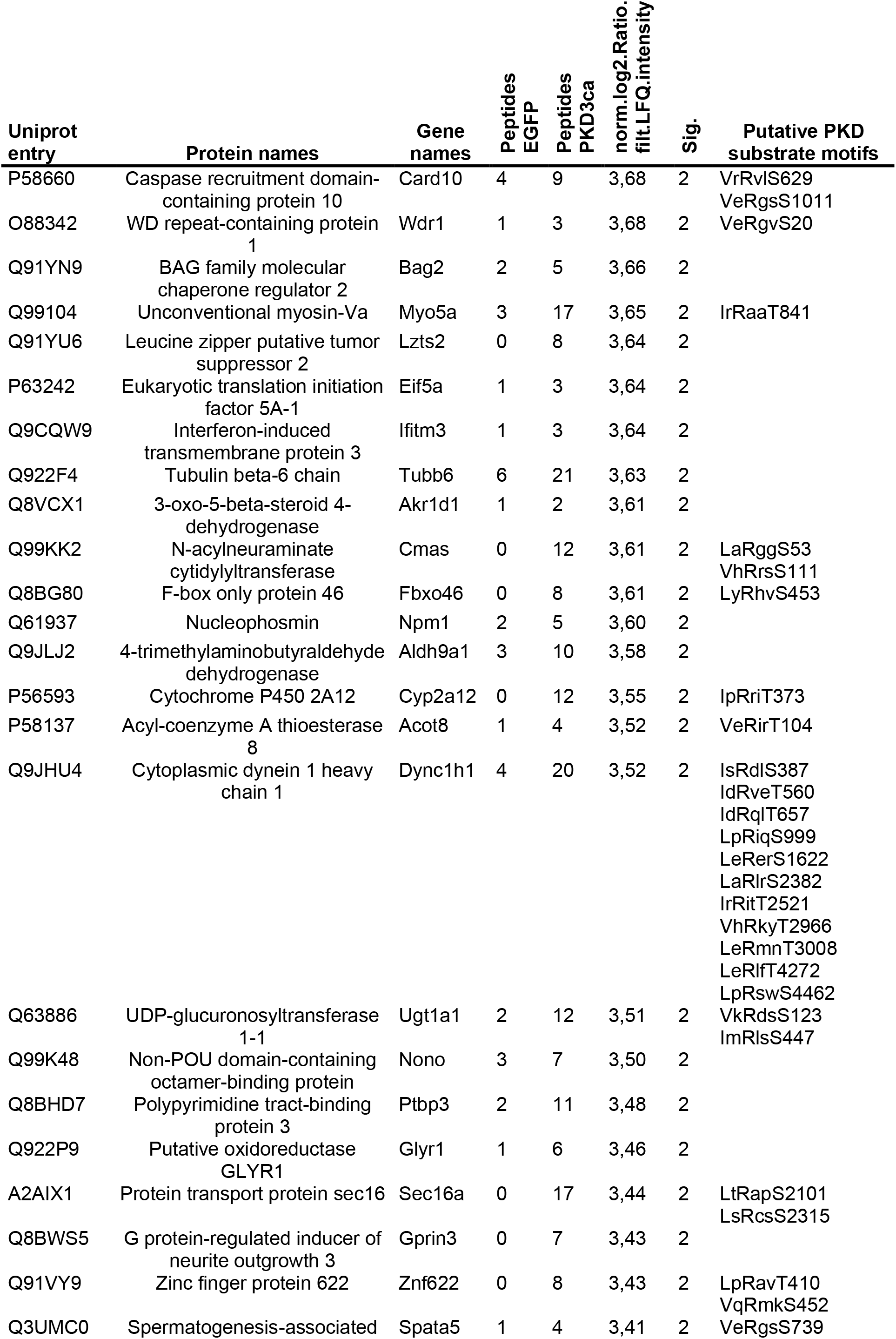
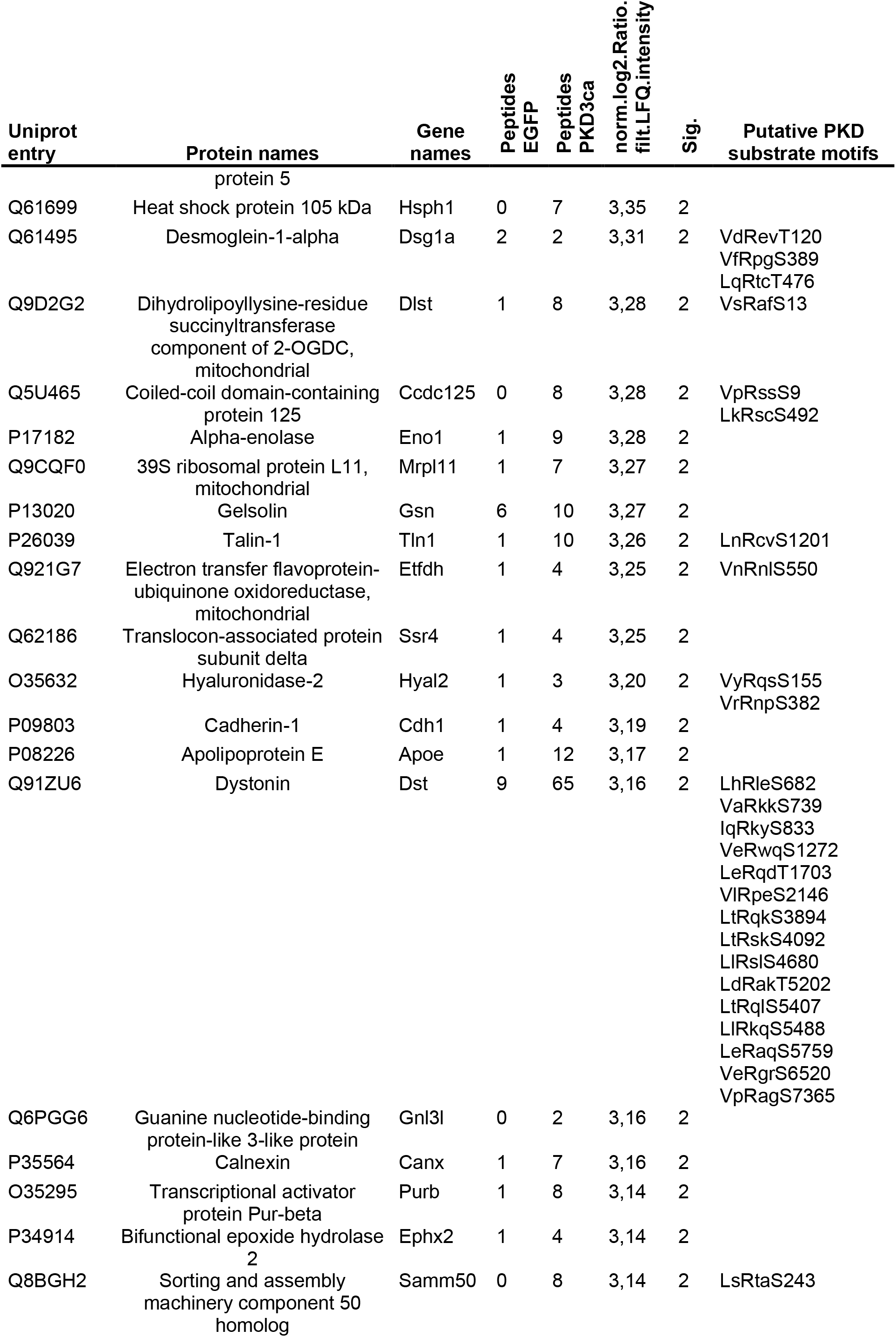
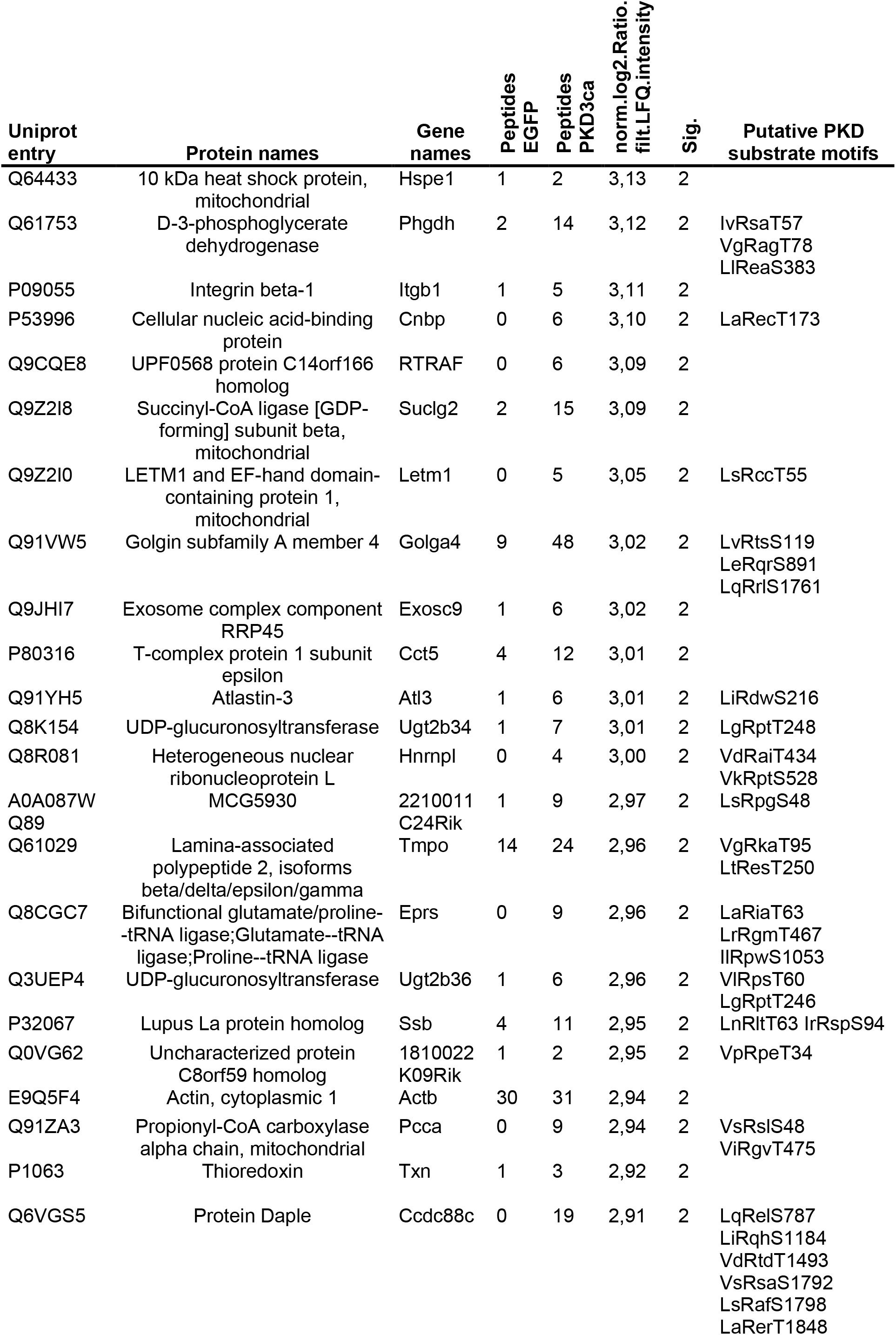
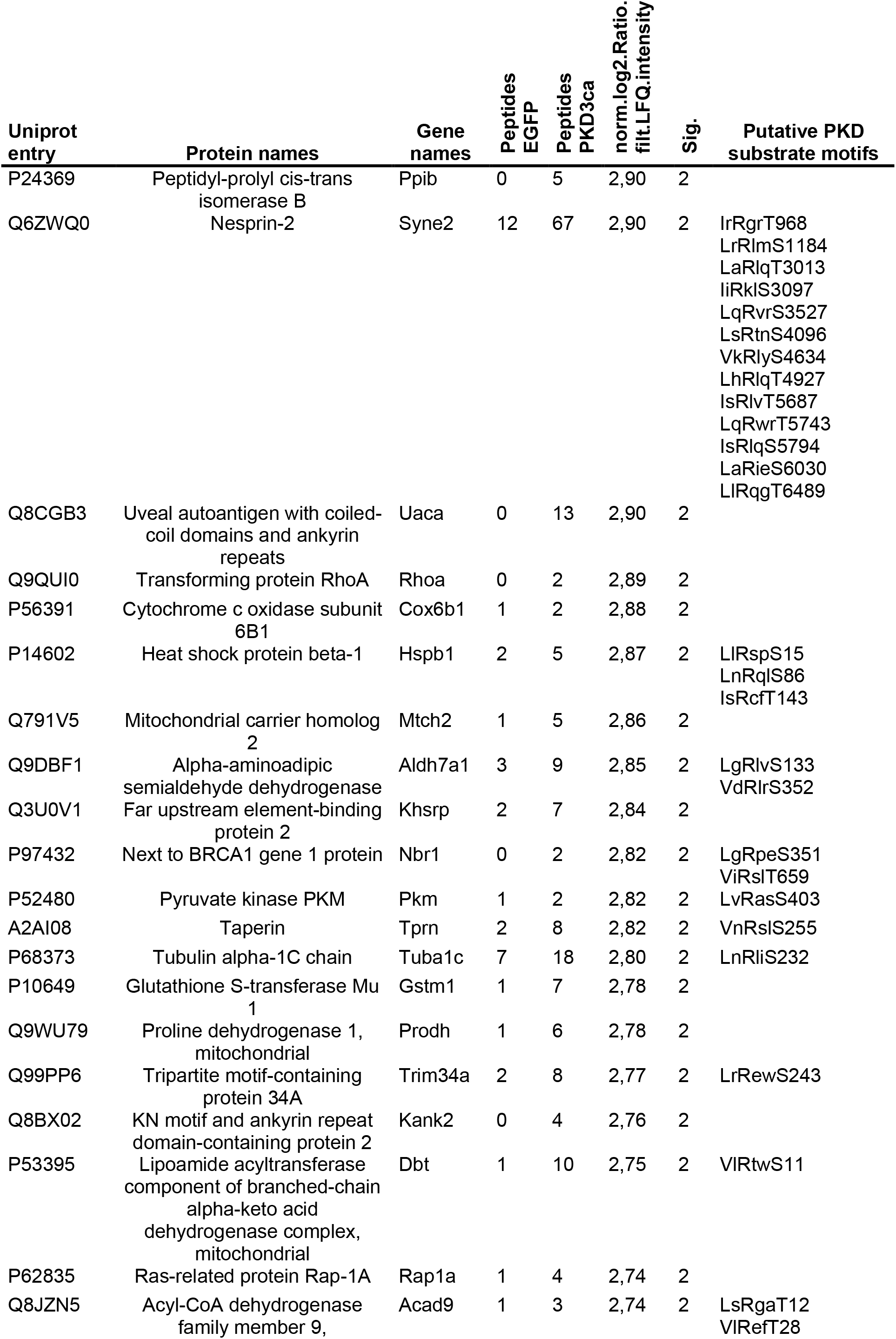
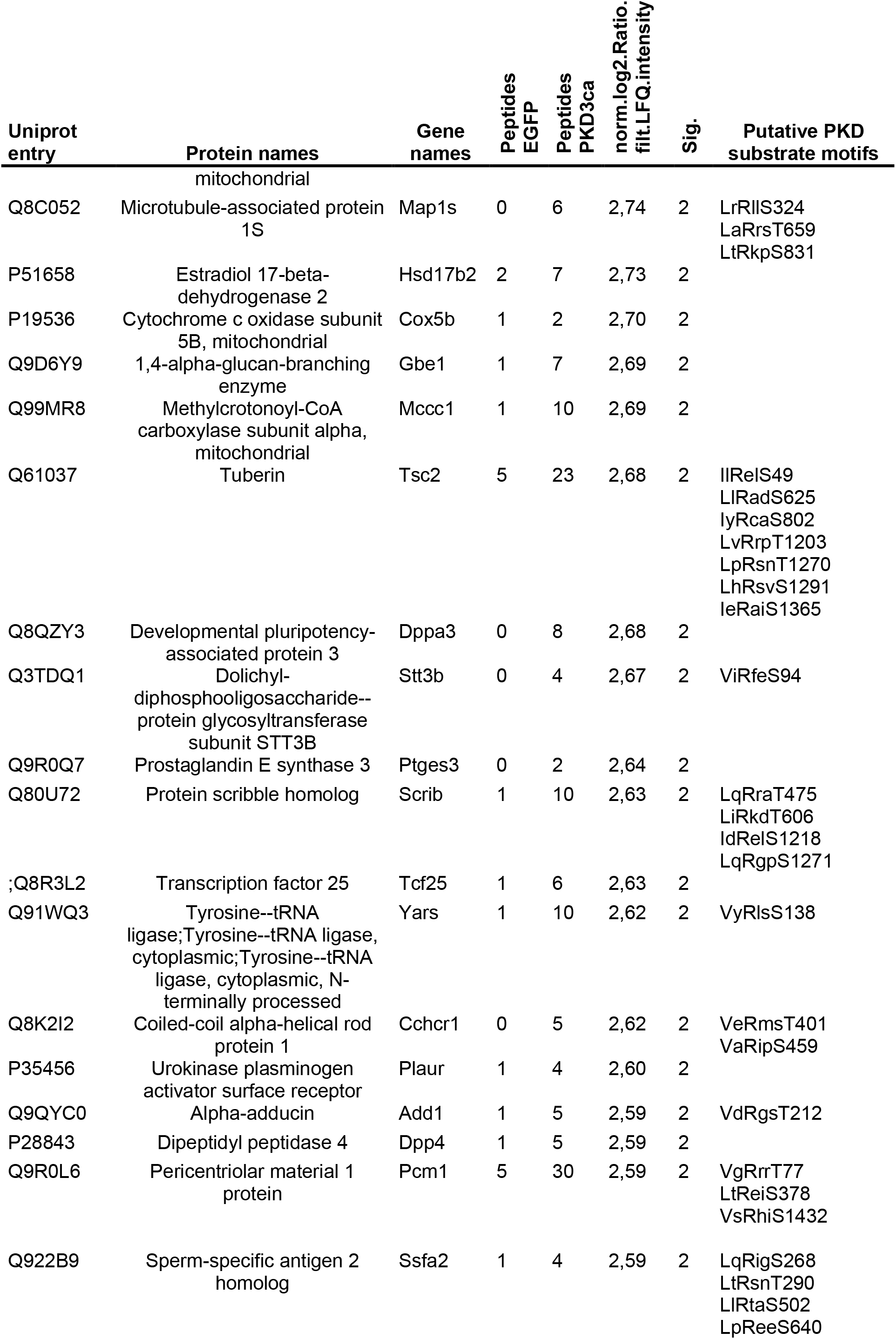
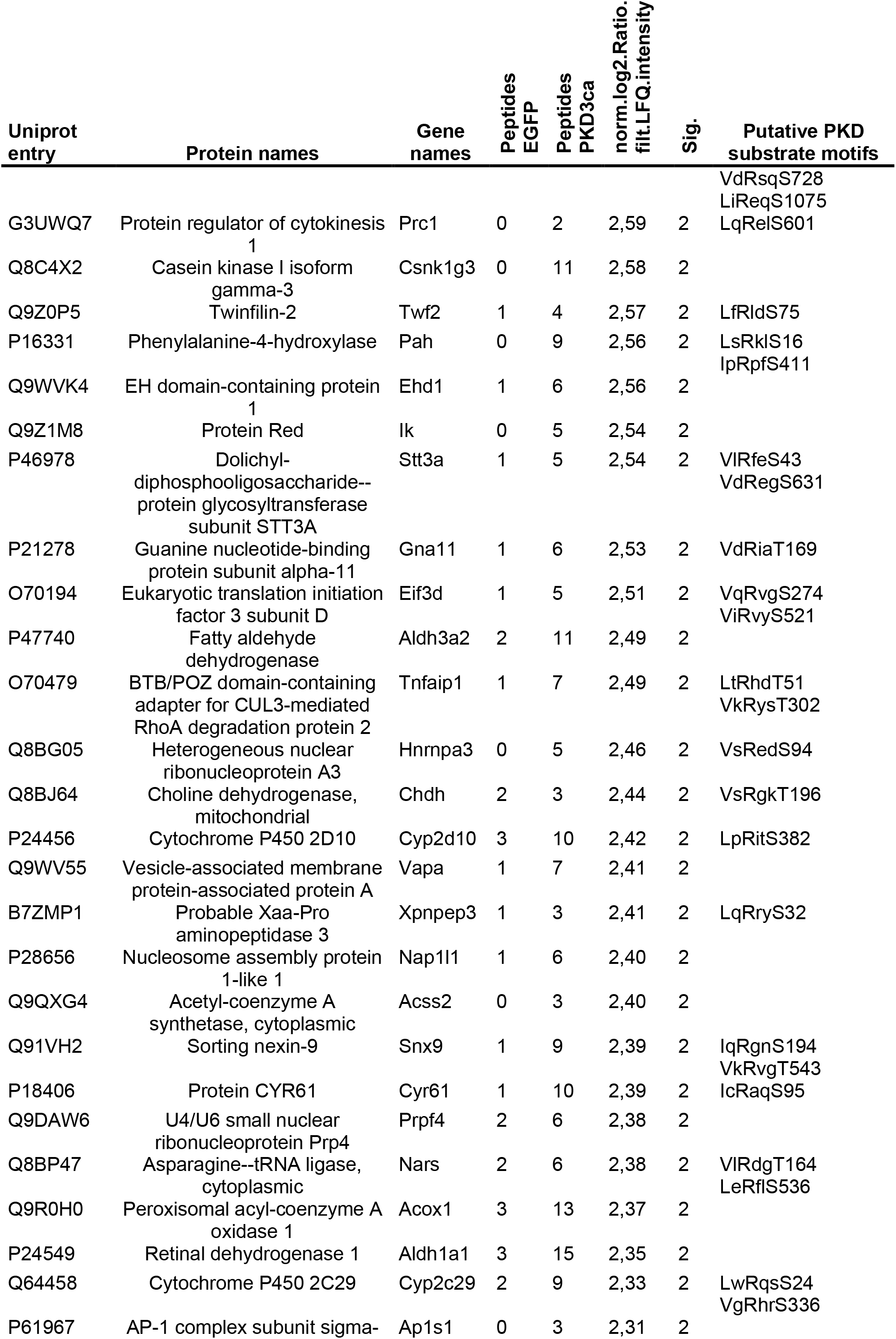
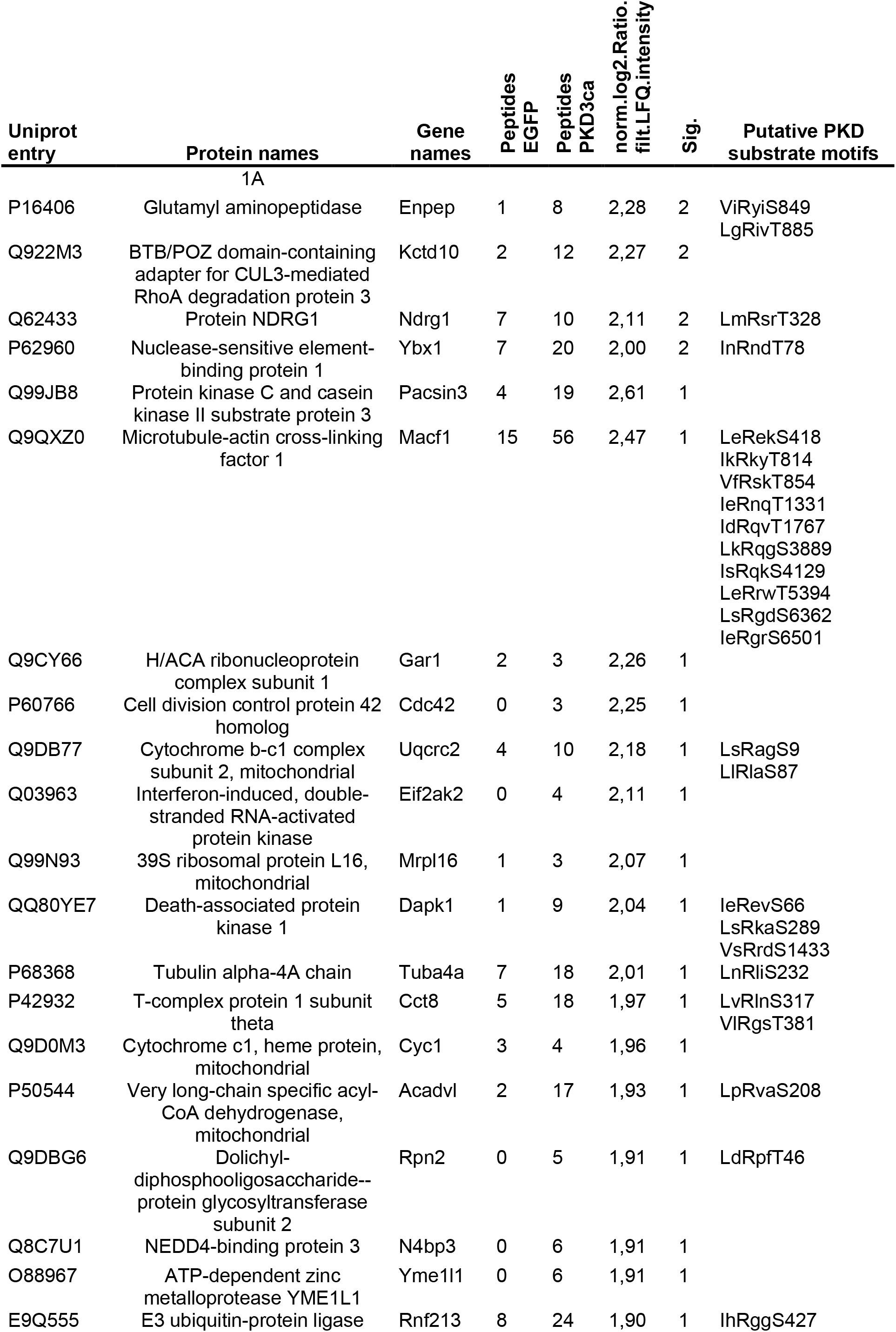
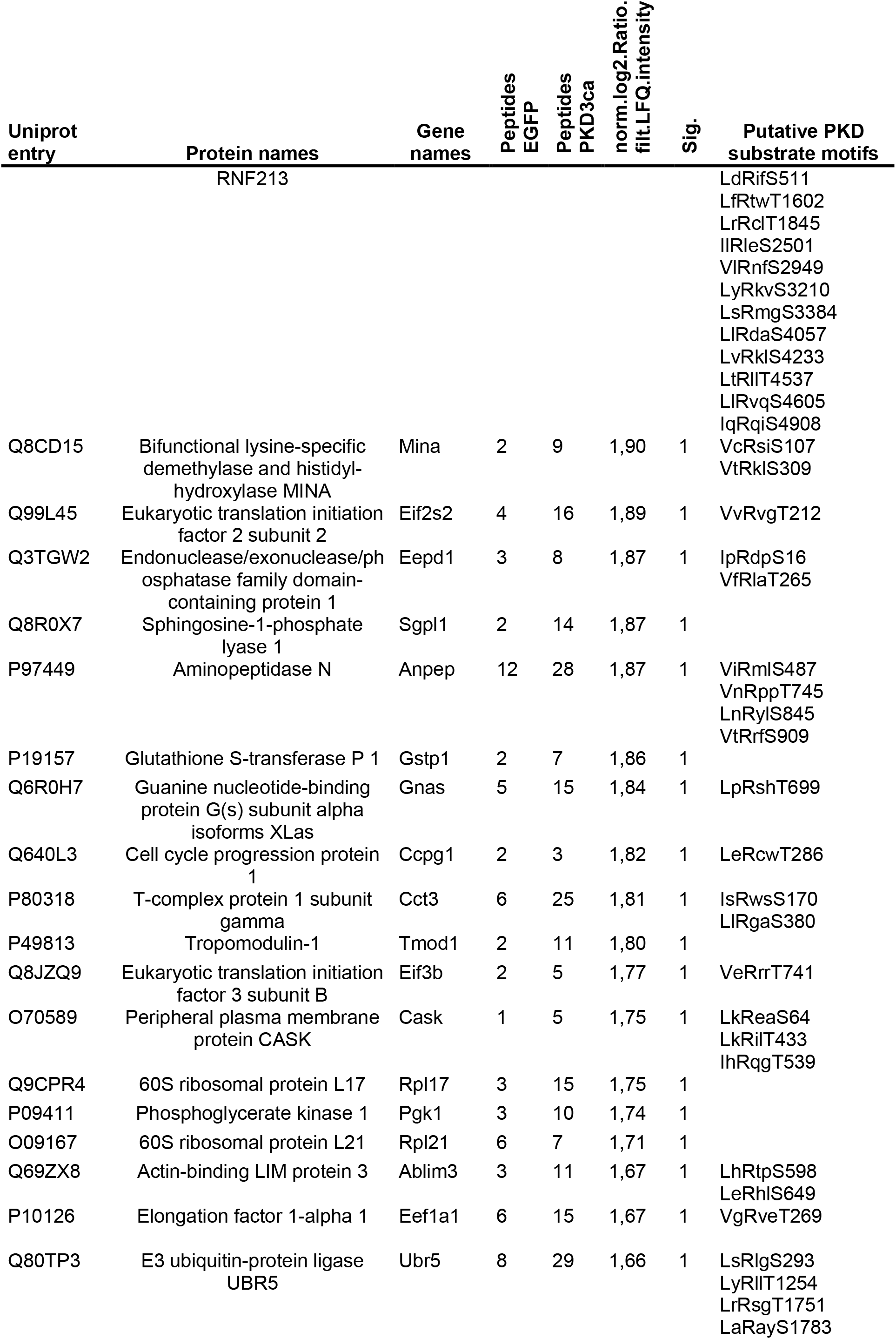
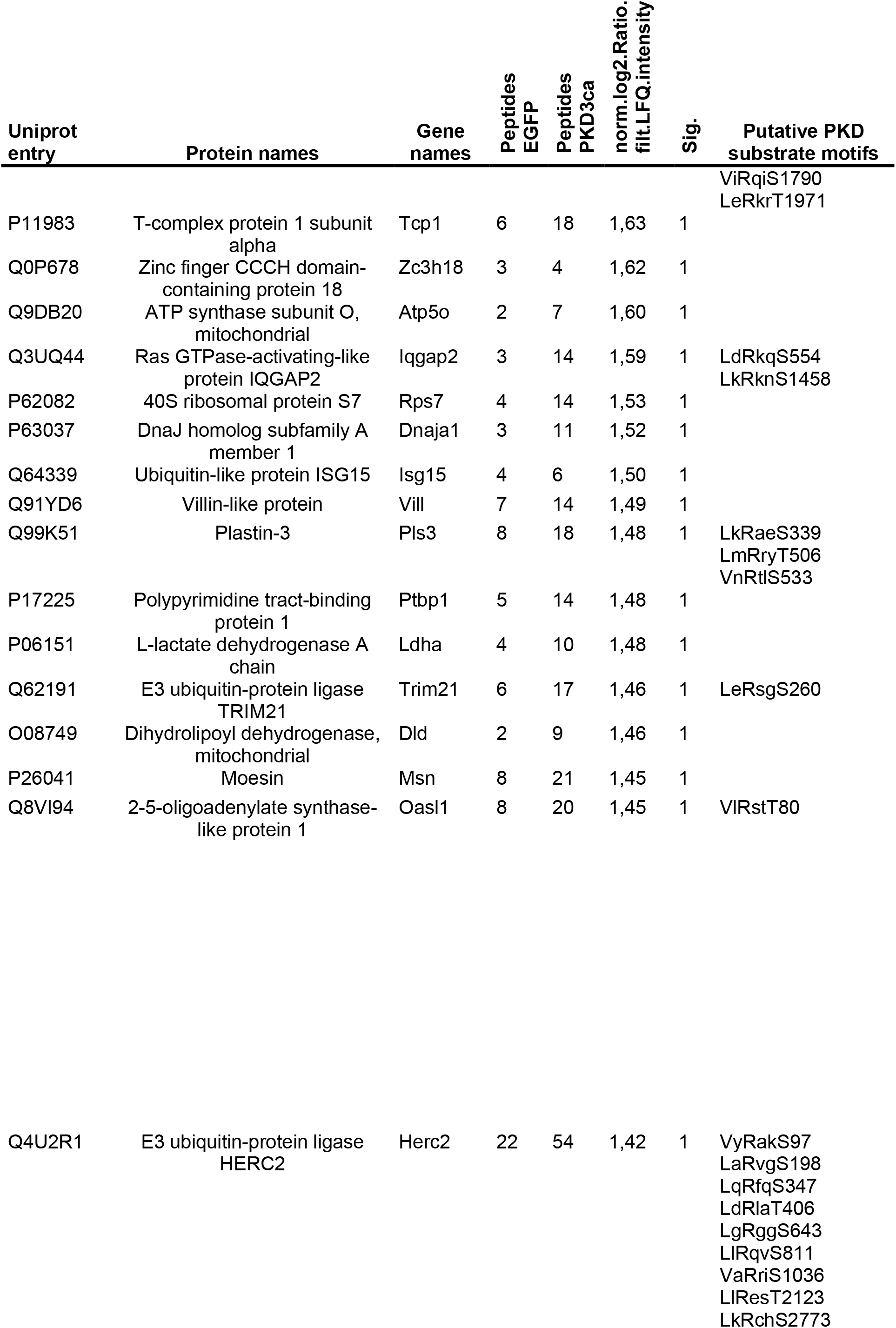
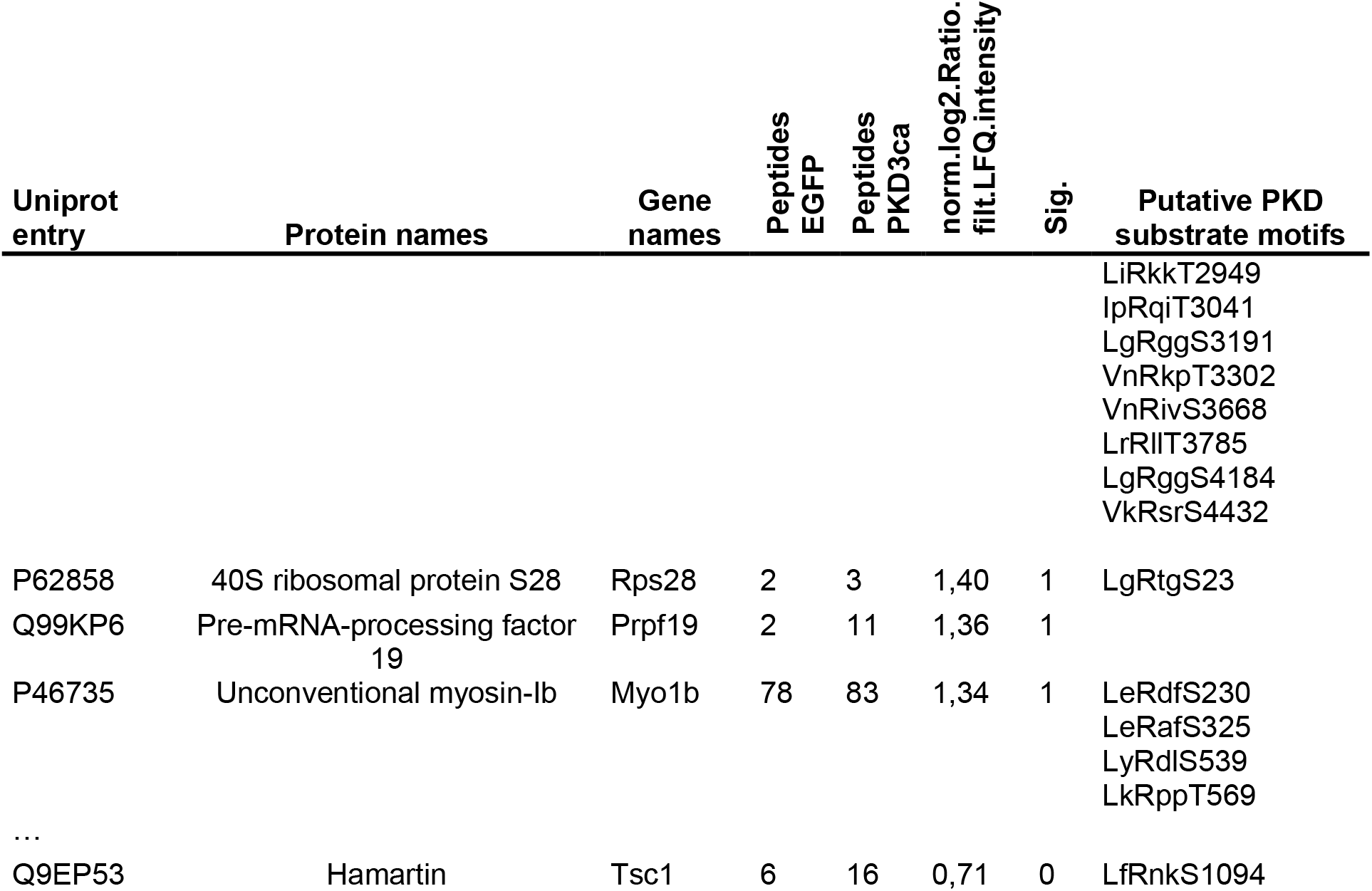
Proteins identified by MS from IP with Rxx[S*/T*] antibody

